# Rapid measurement and statistical ranking of leaf drought tolerance capacity in cotton

**DOI:** 10.1101/2023.08.30.555618

**Authors:** Xuejun Dong, Dale A. Mott, Jhanvi Garg, Quan Zhou, John Sunoj V. S., Benjamin M. McKnight

**Affiliations:** Texas A&M AgriLife Research, Uvalde, TX 78801, USA; Texas A&M AgriLife Extension Service, College Station, TX 77843, USA; Department of Statistics, Texas A&M University, College Station, Texas 77843, USA

**Keywords:** Bayesian hierarchical linear model, drought tolerance ranking, fiber quality, leaf dry matter content, leaf osmotic potential at full turgor, lint yield

## Abstract

Recent progress in ecological remote sensing calls for a more rapid measurement and a closer assessment of crop drought tolerance traits under field conditions. This study addresses three main questions: (1) If leaf dry matter content (LDMC) is equally effective in indicating cotton drought tolerance as leaf osmotic potential at full turgor (*π*_*o*_); (2) if drought tolerance is inversely related to fiber yield/quality in line with the leaf economics spectrum; and (3) if a reliable statistical model can be developed to rank cotton drought tolerance. The values of *π*^*o*^, along with those of LDMC, of 2736 leaves obtained from cotton variety trials conducted during 2020-2022 in both dryland and irrigated regimes were measured using osmometry. The relationships between *π*^*o*^ and LDMC, as well as those between traits and lint yield and fiber quality indices, were investigated using regression analysis. A Bayesian hierarchical linear model was developed to rank cotton drought tolerance based on differences (or adjustments) in *π*_*o*_ and LDMC between dryland and irrigated sites. LDMC was not only shown to be an alternate and equally effective drought tolerance trait compared with *π*_*o*_ obtained from the widely accepted osmometry method, its use is also estimated to lead to a tenfold increase in measuring speed. A stronger drought tolerance capacity of the tested cotton varieties correlated with a lower lint yield and quality, which is generally consistent with the prediction of the leaf economics spectrum. The drought tolerance rankings using the Bayesian hierarchical model help divide the selected 17 cotton varieties into three groups: (a) more-drought tolerant, (b) less-drought tolerant, and (c) intermediate. The ranking results are interpreted using field-measured data of root distribution and diurnal leaf gas exchange from selected cotton varieties. Our work provides new opportunities for a more rapid measurement and an unambiguous ranking of drought tolerance capacity for crop genotypes under various management regimes.

## Introduction

Several decades since the introduction of thermodynamics to describe water potential of plant cells (1), the advent of the pressure-chamber for field measurement of leaf water potentials (2), and the development of the Soil-Plant-Atmosphere Continuum (SPAC) framework (3), the field of plant water relations research has matured into a paradigm with a set of well-tested concepts and models (4–8). In ecological research, however, it is becoming increasingly necessary to develop new methods for rapid measurement of plant water relations traits, which to a large extent has been motivated by the recent progress in remote and proximal detection of plant water status (9–12) and its potential for applications leading to crop improvements (13). A difficult issue arises when the high-throughput sensor estimates of leaf water status need to be calibrated against the measured values, because the latter of which are usually more time-consuming to measure using the pressure-volume techniques (14–16). This issue has recently been mitigated significantly when (17) used the osmotic potential at full turgor (*π*_*o*_) measured relatively rapidly with a vapor-pressure osmometer to predict leaf water potential at turgor loss point (*ψ*TLP), which is evidenced by its wide use (18–26). Nonetheless, the method is still not sufficiently fast enough to measure a large number of leaf samples within a short time period, since the samples need to go through the freezing-thawing cycle and it usually takes 10-15 minutes to measure one sample.

Adjustments in cellular solute concentration (osmotic) and/or cell-wall rigidity (elastic) can be critical for plants to cope with water deficit stress (27–29). Osmotic potential may play a more important role under some circumstances (30, 31), while in others, changes in cell-wall elasticity may be more critical for plants to continue to absorb water in drying soils (32, 33). However, a significant increase in solute concentration in a drying plant leaf cell may lead to its rupture/tissue damage upon rehydration, if not accompanied with an adequate level of cell wall rigidity (28). Thus, osmotic potential and cell-wall elasticity (*ϵ*, or the change in water potential per unit change in volume) in plant leaf cells should co-vary to confer drought tolerance (24, 28, 34, 35). This suggests the possibility of using *ϵ* as a proxy of *π*_*o*_. Yet, *ϵ* is typically estimated through the slow-process of the pressure-volume method and is infeasible to measure for a large number of leaves. Leaf tissue density strongly correlates with *ϵ* (33, 36), but its determination typically involves measuring plant tissue volume, which is not only slow but also prone to errors (37, 38). A good candidate could be leaf dry matter content (LDMC), which is robust and rapid to measure, and may predict *ϵ*, because it has been shown to indicate leaf density and water status (24, 39–41). Based on the above reasoning, we suggest testing the possibility of ‘elevating’ the status of the long-used drought response trait, LDMC, to the same level as *π*_*o*_. We anticipate that, if this is verified through experimental data, it would be feasible to use LDMC to screen a larger number of crop genotypes, as compared to that allowable with the current osmometry method of (17). This is our first interest and is tested in cotton (*Gossypium hirsutum*) in the present work.

We are also interested in linking leaf drought tolerance with crop yield and quality. Consistent with the prediction of the ‘fast-slow’ leaf economics spectrum (LES) (42, 43), it has been shown that a higher capacity of drought tolerance is correlated with a lower photosynthetic potential (33, 44). However, whether a higher drought tolerance consistently results in a lower agronomic yield or quality is not adequately documented, since yield data are typically not available in drought tolerance studies (20). With unusually high temperatures and prolonged drought stress occurring frequently in many of the world’s arid and semi-arid regions, the updated information on drought tolerance capacities of the current crop cultivars is much needed for farmers and crop scientists alike. Specifically, the present work will mainly be based on data of *π*_*o*_ and LDMC for multiple cotton varieties collected over three years.

Our third interest is in ranking drought tolerance capacity among cotton varieties. The main novelty of our approach is that the two response variables LDMC and *π*_*o*_ are used simultaneously so that we can obtain a more precise ranking of the cotton varieties with a relatively small sample size. This is achieved by introducing a latent ranking variable into our linear model and using Bayesian hierarchical modeling to ensure that the ranking of varieties based on LDMC is the same as the ranking based on *π*_*o*_. Further, since this multi-year trial was co-sponsored by industry and the specific varieties to be tested at a specific location/year were not totally decided by researchers, the experimental design is highly unbalanced, posing challenges to the use of conventional statistical methods such as ANOVA (26, 45). But the proposed Bayesian model is highly flexible and can be straightforwardly applied to the unbalanced data we have. We use Markov chain Monte Carlo sampling to find the posterior distribution of the ranking, which illustrates to what extent the observed differences in leaf drought tolerance capacity among cotton varieties is credibly different.

Finally, we are interested in interpreting cotton leaf drought tolerance in terms of water use strategy. Under uncertain water supply, an effective stomatal control of water loss is critical to plant survival and growth (46), and it requires an integration of stomatal conductance (*g*_*s*_) with multiple environmental variables (47). Recent progress (48) provides a mechanistic interpretation of the dynamics of *g*_*s*_ that is compatible with both earlier empirical (47, 49) and theoretical work (50) of stomatal optimization. We want to test if the more drought tolerant cotton varieties also have a more conservative strategy of water use in leaf gas exchange. Another key factor governing plant water use is rooting depth (51). It has been observed in woody species—cotton is also woody species, but is managed as an annual crop—that a shallower root system corresponded to a suite of more drought tolerant traits (32, 52, 53). Thus, we, similar to (35), want to ask if cotton varieties with a higher drought tolerance capacity (namely, more negative *π*_*o*_, or larger LDMC, or larger adjustment in *π*_*o*_ or LDMC) have a shallower root system. The objectives of the present work are to:

1. Compare leaf osmotic potential at full turgor (*π*_*o*_) and leaf dry matter content (LDMC) as drought tolerance traits in cotton.
2. Interpret the linkage between measured leaf drought tolerance traits and cotton lint yield and fiber quality indices.
3. Conduct statistical ranking of drought tolerance capacity for multiple cotton varieties.
4. Interpret drought tolerance responses of selected cotton varieties based on measured water use traits of leaf gas exchange and rooting depth.

## Materials and Methods

### A. Description of the field sites, experimental design and crop management

The study was conducted in conjunction with Texas A&M AgriLife Extension Replicated Agronomic Cotton Evaluation (RACE) trial. Six cotton fields from four locations in southwest to central Texas, USA (i.e., Crystal City, Uvalde, Lytle, and Taylor) were selected for field sampling (see Supplementary Figure A1). The field in Taylor was under dryland production, and the remaining fields were under irrigated management from 2020 to 2022. About 71 cotton varieties were used in the study (See Supplementary Material A for detail). Most of the varieties were from the RACE trial planted in Lytle and Taylor. Sixteen of them were also planted at the Uvalde Research Center. In producers’ fields in Uvalde and Crystal City, only one or two varieties were used. All the cotton fields were planted from mid- to late April, except the Uvalde Research Center field in 2020, which was planted on May 5, 2020. Management of the cotton crop at all sites followed the recommended practice for the region. Leaf area index (LAI) from all fields was measured once every two weeks after May 15, 2020. The measurement was made using LI-2000 Canopy Analyzer (LI-COR Inc., Lincoln, NE, USA; see Figure 1). At each field (except the one in Taylor), the amounts of irrigation applied and rainfall received during the main growing season were measured using 2 standard rain gauges (All Weather Rain Gauge, Scientific Sales, Inc. Lawrenceville, NJ, USA). Later, the irrigation amount was estimated by subtracting the amount of daily rainfall (obtained from the 1-km grid weather data product available at https://daymet.ornl.gov) from the total amounts measured using the rain gauges.

**Fig. 1.**
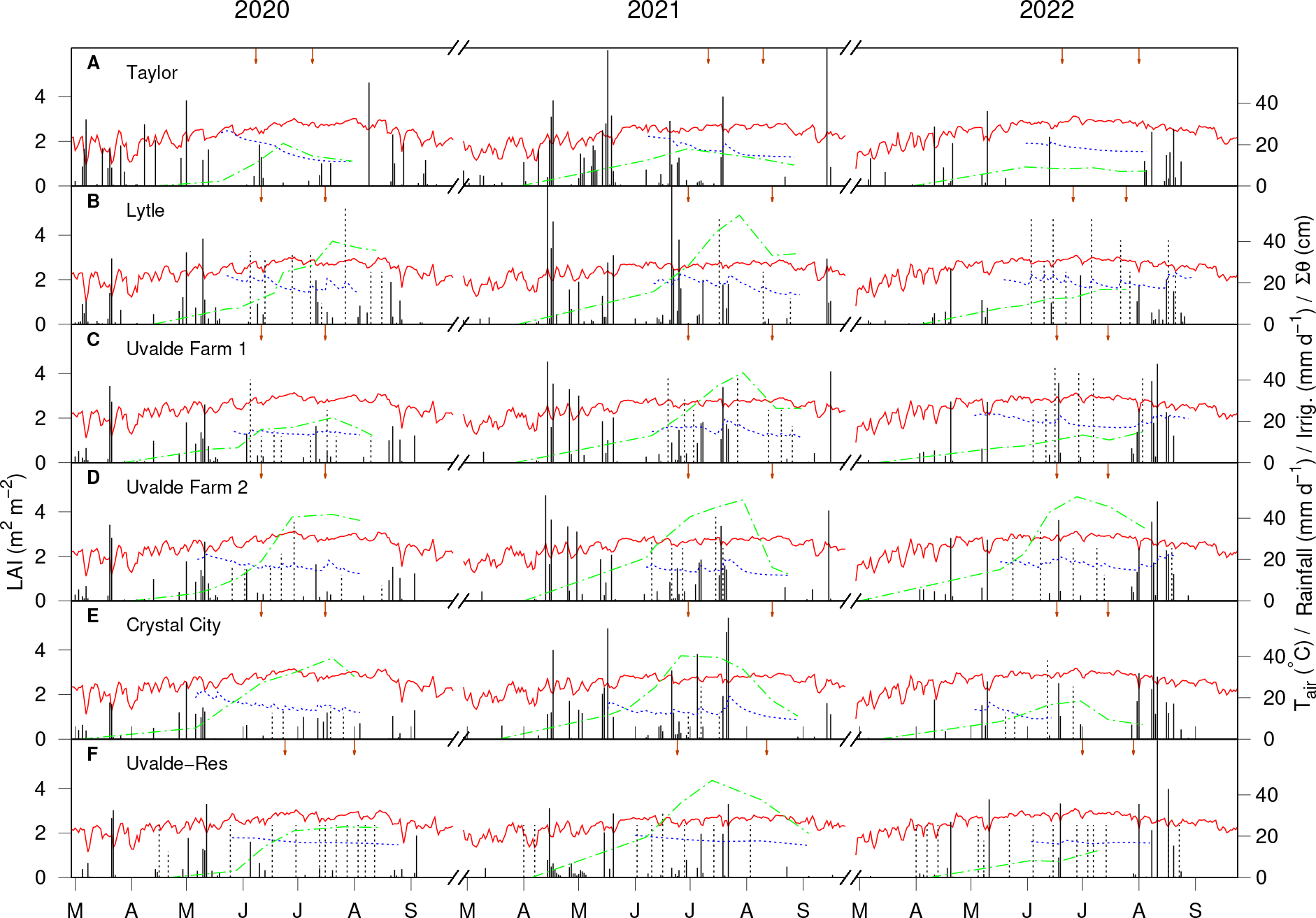
Daily weather data, including average air temperature (°C, solid lines), rainfall (mm day^−1^, solid impulses), and irrigation (mm day^−1^, dotted impulses), soil water storage in top 60 cm soil depth (Σ*θ* in cm, dotted lines) and leaf area index (m2 m^−2^, dash-dotted lines) during the growing seasons of 2020 through 2022 at Taylor (**A**), Lytle (**B**), Uvalde Farm # 1 (**C**), Uvalde Farm #2 (**D**), Crystal City (**E**), and Uvalde Research Center (**F**). Each year at each location, the plotted daily data values range from March 13 to October 8, and each of the horizontal tic marks indicate day 15 of a consecutive month from March (M) to September (S). The two down arrows located in a specific growing season and field site indicate the actual dates when the cotton leaf samples were collected, roughly in early and peak bloom stages, respectively. Weather data at the Uvalde Research Center site were measured from an on-site weather station, while data from all other sites were obtained from the 1-km grid weather data product available at https://daymet.ornl.gov (54). Data of leaf area index and soil water storage were measured at each site.

### B. Measurement of leaf drought tolerance traits

Over the three years, a total of 2736 leaves from 71 cotton varieties (names listed in Supplementary Tables A1-A3) were collected to measure drought tolerance traits. Leaf samples were collected twice: first at the early bloom and then at the peak bloom stage (arrows in Figure 1). On each sampling, three leaves per variety/treatment were cut off at the base of the petioles from the 4th node of three representative plants. Then the petioles of the leaves were immediately submerged in a 2-cm deep of distilled water in a bucket. On the same day, the collected leaves were stored in a laboratory overnight with the top of the bucket covered with aluminum foil to maintain a high humidity condition in the interior of the bucket. The next day, water droplets from the surfaces of the leaf were gently blotted using tissue paper and immediately prepared for physiological measurements. One 8-mm leaf disc was punched off of each of the sampled leaves, wrapped in aluminum foil, immediately frozen in liquid nitrogen, and then stored at a -80 °C freezer awaiting osmotic potential measurement. In punching the leaf discs, the main veins were avoided, following (17). Immediately after the processing of the punched leaf disc, the leaf was measured for saturate mass and area, then dried at 65 °C to measure dry mass. Leaf osmotic potential was measured using a 5520 VAPRO Vapor Pressure Osmometer (Wescor, Inc., USA), following (17). Leaf dry matter content (LDMC) was determined as the ratio of dry mass and saturate mass. In this study, the measured leaf osmotic potential at full turgor (*π*_*o*_) was used as a proxy of leaf water potential at leaf turgor loss point (*ψ*TLP), and the measured LDMC was used to indicate the cell wall elastic modulus (*ϵ*).

### C. Measurement of soil water content

At the Uvalde Research Center, twelve access tubes were installed (two in each of the six plots) in May 2020, 2021, and 2022, and soil water content was measured using a CPN 503 Elite Hydroprobe (InstroTek, Inc., Research Triangle Park, NC, USA) from 0-120 cm depth at intervals of 10, 20, 40, 60, 80, 100, and 120 cm. The neutron probe was calibrated using the twopoint method as recommended by the manufacturer. Soil water data were collected six to nine times from early June to early September. Soil water content in the remaining fields was measured at 15, 30 and 61 cm depths using a set of soil moisture sensors (5-TM, GS-1, GS-3, or Tero-12, METER Group, Inc., Pullman, WA, USA) connected to EM-50 dataloggers (METER Group, Inc., Pullman, WA, USA) from early May to early September. Daily averaged soil water content was calculated based on the 60-min raw data.

### D. Measurement of leaf gas exchange and water status

Leaf gas exchange rates were measured on cotton plants that were sown on April 22, 2022 in 0.76-m row spacing in a center pivot field under full- and deficit irrigation regime, receiving 300 mm and 254 mm water input, respectively, during the crop season (rain + irrigation). Leaf gas exchange parameters, including net photosynthetic rate (*A*), stomatal conductance (*g*_*s*_), and intercellular CO_2_ concentration (*C*_*i*_), were measured using a unit of LI-6400XT Portable Photosynthesis System (LI-COR Bioscience, Lincoln, NE, USA). The intrinsic water use efficiency (iWUE) and carboxylation efficiency (CE) were calculated as the ratio of *A/g*_*s*_ and *A/C*_*i*_, respectively, and instantaneous WUE was calculated as *A/E*, where *ϵ* is transpiration rate (mmol H_2_O m^−2^*s*^−1^) (55). The measurement was made on July 20, 2022 (clear day) at 8 am, 11 am, and 3 pm on fully expanded sunlit leaves from the upper canopy of three randomly-chosen plants, each from varieties DP 2020, DP 1725, and PHY 400. The reference CO_2_ concentration of the photosynthesis system was set to 400 *μ*mol mol^−1^ using a 6400-01 CO_2_ mixer installed with a 12-g pure CO_2_ cylinder. The system flow rate was set to 500 *μ*mol s^−1^ and the chamber cuvette fan at high speed. Before clamping a leaf into the chamber, the ambient photosynthetically active radiation (PAR) incident on the leaf was measured using an external quantum sensor (LI-190R) attached to the leaf chamber and the same PAR was provided to the leaf inside the leaf chamber with a 6400-02B LED light source. The measurements were made under ambient relative humidity and temperature, and on a fixed leaf area of 2 × 3 = 6 cm^2^. The sensor head of the photosynthesis system was placed in shade during the waiting time between measurements. Leaf chlorophyll fluorescence transients, canopy temperature (*T*_*C*_), and leaf water potential (*ψ*_*L*_) were also measured. Each time, a minimum of four chlorophyll fluorescence and three *ψ*_*L*_ measurements was recorded from each variety. Maximum photochemical efficiency of photosystem II (PSII; *F*_*v*_*/F*_*m*_) and thylakoid membrane damage (*F*_0_*/F*_*m*_) were measured from similar leaves as used for gas exchange measurement after 30 min of dark adaptation using a chlorophyll fluorometer (OS1p; OptiSciences, Hudson, NH, USA). *F*_*v*_*/F*_*m*_ was calculated as *F*_*v*_*/F*_*m*_ = (*F*_*m*_ −*F*_0_)*/F*_*m*_, where *F*_0_ and *F*_*m*_ are the minimum and maximum fluorescence, respectively, in the dark-adapted state. A light pulse intensity of 3000 mmol m^−2^ s^−1^ with a duration of 3 sec was applied to the leaf to generate *F*_*m*_. Leaf water potential (*ψ*_*L*_) was measured using a pressure chamber (Model 615; PMS Instrument Company, OR, USA), with the high pressure provided by compressed pure nitrogen gas. Canopy temperature (*T*_*C*_) was measured using a hand-held infrared radiometer (MI-210, Apogee Instruments Inc, UT, USA).

### E. Measurement of vertical root distribution for selected cotton varieties

Vertical root distribution was measured by digging the soil in the vicinity (10-cm away from the main stem) of selected cotton plants at the Taylor site on July 7, 2023. This was at the early bloom stage with presumably the greatest rooting depth (56). An auger with a 12-cm diameter corer was used. The soil was sampled at 10-cm intervals within the top 100-cm depth, and the sampled soil was collected in 3.78 liter ziploc bags, which were later stored in a -10°C freezer the same day of sampling and await further processing. The soil samples were collected only from 12 plots, representing four cotton varieties (DP 2012, NG 4190, PHY 400 and PHY 415) replicated 3 times with a randomized complete block design). The 120 frozen soil samples were then washed and rinsed using a 0.5-mm mesh testing sieve (W. S. Tyler Incorporated, Mentor, OH, USA) to collect roots in 20-ml scintillation vials filled with 20% ethanol. The roots from each of the sample vials were further cleaned to remove foreign objects, spread and floated in a water-filled tray to minimize cluttering and overlapping of rootlets, and scanned into computer to create a digital image, which was then analyzed using the WinRHIZO software ver. 2020 (Regent Instruments Inc., Canada) to calculate total root length (cm). Root length density (*L*_*d*_) of each of the samples (volume ≈ 1130 cm^3^) was expressed as cm cm^−3^.

### F. Measurement of cotton lint yield and fiber quality indices

From 2020 to 2022, cotton yield was measured by harvesting the entire plots at Lytle (each plot 6 rows 367 m in 0.91-m row spacing) and Taylor (6 rows 320 m in 0.97-m row spacing). The Lytle site was harvested with a cotton picker whereas the Taylor site with a cotton stripper. Sub-samples of the plots were ginned on a 20 saw Centennial Gin for turnout and lint samples were then used to obtain fiber quality at Fiber & Biopolymer Research Institute at Texas Tech University. The fiber quality indices include: micronaire, length, strength, and uniformity. Cotton yields and fiber quality data from farmers’ fields at Uvalde and Crystal City were not available since the farmers declined to provide yield data.

### G. Data analysis

#### G.1. Estimation of soil water stress index

For each field site and at each leaf sampling period, the average value of the site-specific soil water stress index (SWSI) was calculated based on measured soil water content described in Section C:

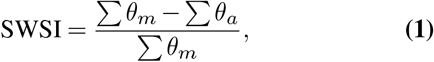

where Σ *θ*_*m*_ and Σ *θ*_*a*_ are sums of maximum and actual available water, respectively, in the 60-cm soil profile:

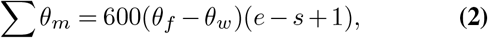

where *θ*_*f*_ and *θ*_*w*_ represent water content at field capacity and wilting point, respectively, and *e* and *s* represent the ending day number and starting day number for a time period.

The value of Σ *θ*_*a*_ for a particular site/environment for a particular time period was calculated by adding the summed available water at the three soil depth intervals (namely at 150 mm, 150 mm, and 300 mm, respectively):

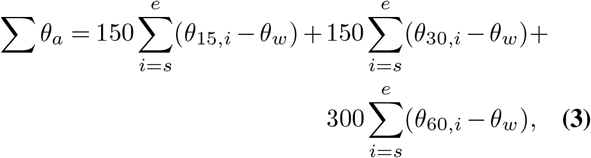

where *θ*_15,*i*_, *θ*_30,*i*_ and *θ*_60,*i*_ are average values of volumetric water contents on the *i*th day at 15 cm, 30 cm. and 60 cm soil depth, respectively. The reason for using the summations is to get an estimate of the average soil water deficit that cotton plants would have to experience during a certain period of time covering the first or second sampling of the cotton leaves in a particular field.

#### G.2. Analysis of the relationships between leaf osmotic potential at full turgor and LDMC and their linkages with cotton lint yield and fiber quality indices

Linear regression analysis was used to describe the relationships between *π*_*o*_ and LDMC measured at the early and peak bloom stages for leaf samples collected from irrigated and dryland sites separately. Where appropriate the statistical differences of the regression lines were compared using ANOVA with GraphPad Prism 6 (Version 6.07 for Windows, 2015, GraphPad Software, San Diego, CA, USA), along with the Tukey’s multiple comparisons test (*α* = 0.05).

#### G.3. Statistical ranking of cotton varieties for drought tolerance

We rank the cotton varieties based on the adjusted value of *π*_*o*_ or LDMC, defined as the difference between the measurement taken at the dryland (Taylor) and that at the irrigated plot (Lytle). So, only those varieties which were used both in the dryland and irrigated sites in the same season were chosen. This results in a data set consisting of *p* = 17 varieties in *m* = 6 different seasons; see Supplementary Table C1. To rank the drought tolerance capacity of different varieties using LDMC and *π*_*o*_ jointly, we build a Bayesian hierarchical linear model which can efficiently handle the missing data and integrate the information of LDMC and *π*_*o*_ measurements.

Denote the set of *p* cotton varieties shown in Supplementary Table C2 by 𝒱 and the set of *m* seasons in the same Table by 𝒮. In total, we have *n* = 280 replicates (which henceforth will also be referred to as ‘observations’). For the *k*-th observation (*k* = 1, 2, …, *n*), let *Y* ^D^(*k*) denote the standardized LDMC measurement, *Y* ^*π*^(*k*) denote the standardized *negative π*_*o*_ measurement, vat(*k*) ∈ 𝒱 denote the cotton variety and sea(*k*) ∈ 𝒮 denote the season. Further, let env(*k*) denote the environmental condition of the *k*-th observation, with env(*k*) = 1 for the dryland site and env(*k*) = 0 for the irrigated site. We assume that

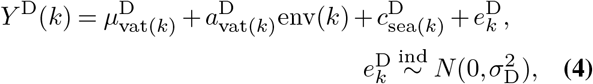

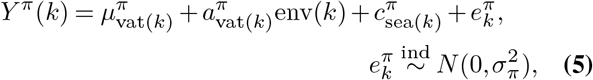

where 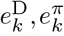 are independent Gaussian errors, 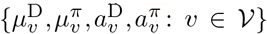 and 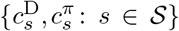 are unknown parameters of interest, and 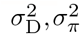 are unknown hyperparameters. For ease of notation, we will also use ***μ***^D^ to denote the vector that contains *p* elements 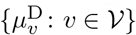, assuming an implicit one-to-one mapping from 𝒱 to {1, 2, …, *p*} ; ***μ***^*π*^, ***a***^D^, ***a***^*π*^, ***c***^D^, ***c***^*π*^ are defined similarly, but note that ***μ***^D^, ***μ***^*π*^, ***a***^D^, ***a***^*π*^ ∈ ℝ^*p*^ while ***c***^D^, ***c***^*π*^ ∈ ℝ^*m*^. The parameter ***a***^D^ (resp. ***a***^*π*^) characterizes the difference in LDMC (resp. *π*_*o*_) measurement for the same cotton variety under dryland and irrigated conditions. If the leaf of a cotton variety *v* has better drought tolerance, then both 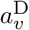 and 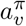 tend to be larger. Since we expect that the magnitude of both LDMC and *π*_*o*_ should increase when the same cotton variety is moved from an irrigated site to a dryland one, we will impose the constraint that 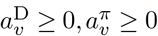 for all *p* varieties. Due to unexpected circumstances, we only observed *Y*^D^ (*k*) but not *Y* ^*π*^(*k*) for *k*’s such that sea(*k*) = 2021_peak, but we will see that this does not cause any issue for the Bayesian inference methodology we propose.

The main idea of Bayesian statistical modeling is to first specify a prior distribution for unknown model parameters and then update it using the data likelihood, which yields the posterior distribution of parameters. Compared to standard Bayesian linear regression models, the key novelty of our method is that our prior forces the drought tolerance parameters 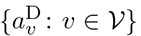 and 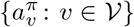 to have the same ranking. This is achieved by introducing a latent variable *τ*, which represents the ordering of *p* varieties, and letting the conditional prior distributions of ***a***^D^ and ***a***^*π*^ given *τ* to be constrained to the same set. We describe our prior in detail in Supplementary Material C.

We use Markov chain Monte Carlo (MCMC) sampling to learn the joint posterior distribution of the parameters *τ*, ***μ***^D^, ***μ***^*π*^, ***a***^D^, ***a***^*π*^, ***c***^D^, ***c***^*π*^, 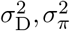. Due to the use of conjugate prior distributions, we are able to build an efficient random-scan Metropolis-within-Gibbs algorithm, detailed in Supplementary Material C. The Metropolis step is used to update the ordering *τ*. We run the sampler for 10^6^ iterations with half of them as burn-in, which only takes a few minutes on a personal laptop. Trace plots suggest that the algorithm has approximately converged after 10^4^ iterations.

#### G.4. Analysis of leaf gas exchange data

Data of diurnal course of leaf gas exchange parameters, along with data of leaf water potential and canopy temperature, were analyzed graphically to visualize the differences among selected cotton varieties at different times during the day when the measurements were made. This was followed by further analysis using the General Linear Model in SPSS Ver. 16 (SPSS Inc., Chicago, IL, USA) considering variety and time of day as fixed factors.

Data of stomatal conductance as a function of photosynthetic rate, vapor pressure deficit, and leaf surface CO_2_ concentration were fitted using a model of (48):

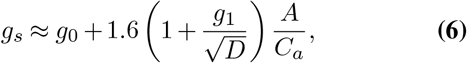

where *A* is net photosynthetic rate (*μ*mol m^−2^s^−1^), *C*_*a*_ is the CO_2_ concentration at the leaf surface (*μ*mol mol^−1^), *D* is the leaf-to-air vapor pressure deficit (kPa), *g*_0_ is minimum stomatal conductance (mol m^−2^s^−1^) and *g*_1_ is proportional to the marginal water cost of carbon gain 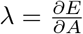 as used by (50) : 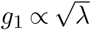 (also see (57)). Since our focus was on parameter *g*_1_, the value of parameter *g*_0_ of Eq. (6) was assume zero (58, 59). The model fitting was done using the Gauss-Newton method via Minitab (Minitab 17.3.1, Minitab, Inc. 2016). The statistical differences in *g*_1_ among cotton varieties were compared using ANOVA with GraphPad Prism 6, along with the Tukey’s multiple comparisons test (*α* = 0.05). Meanwhile, the stomatal behavior described by Eq. (6) was also visualized by plotting *g*_*s*_ against 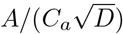.

#### G.5. Estimation of rooting depth for selected cotton varieties

The measured root length density (*L*_*d*_(*z*)) was used through the computer program len.mac of (60) to estimate the soil depth above which 95% of root length is located, which is referred to as ‘rooting depth’ (*z*_95_). Specifically, the normalized root length density is defined according to (61) as

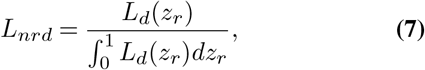

where 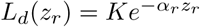 is root length density at relative soil depth *z*_*r*_ = *z/L*_*m*_, *z* and *L*_*m*_ are soil depth (cm) and maximum possible rooting depth (cm), respectively, and *K* and *α*_*r*_ are empirical parameters defining the shape of the *L*_*d*_(*z*_*r*_) curve. Based on our experience for this particular field site, *L*_*m*_ was assumed to be 120 cm for all plots. A one-parameter model of (62) was used to implement Eq. (7):

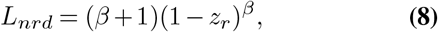

where *β* is an empirical parameter. The values of *z*_95_ for different plots were estimated based on the value of *L*_*m*_ and the respective values of *β* as

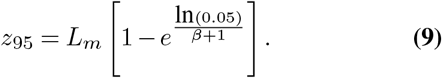

In addition to providing outputs of estimated model parameters *K, α*_*r*_ *β* and the derived variable *z*_95_, len.mac also calculates the ratio of total root lengths at the top and bottom soil profile (*R*_*t*2*b*_). In this site, we chose 40 cm as the division of the top and bottom soil profile. The differences of the model parameters and derived variables among cotton varieties were compared using analysis of means (ANOM) available in Minitab with *α* = 0.10 and *n* = 3.

## Results

### H. Weather, soil water condition and leaf area growth

The amounts of rainfall received during the crop seasons varied greatly, with 1 std. deviation accounting for 29-41% of the 43-year averages (Supplementary Table A4). The severe drought in 2022 affected Taylor and Lytle sites more than the other sites, with seasonal rainfall reduced by 47% and 59%, respectively, from the long-term average. The daily rainfall and irrigation, along soil water storage and LAI, are visualized in Figure 1. The dryland site (Figure 1A) started out with a full soil water profile in early May. It then underwent a nearly monotonous decrease in the next 3 months, except on August 1-2, 2021, when two rain events (57.2 mm of rain) caused a spike in soil water storage. The values of soil water storage in the irrigated sites generally responded to the inputs from rainfall or irrigation, except that at Uvalde Research, in which the response was less clear (Figure 1F). That was because soil water content was measured less frequently (once in 2 weeks) using a neutron probe, as opposed to the automated sensor measurement in other sites. In 2022, the datalogger cable at the Crystal City site was damaged by deer, causing total data loss after June 28 (Figure 1E).

Across three years, LAI in the dryland site (Figure 1A) was no greater than 1.91 m^2^ m^−2^, with the lowest peak LAI (0.82 m^2^ m^−2^) occurring in 2022. LAI values in the irrigated sites generally responded positively to the amount of water inputs, especially before and during the bloom stages. In 2022, the LAI values in Uvalde Research site were low (*<*1.22 m^2^ m^−2^) despite the frequent irrigation (Figure 1F), which was likely caused by weed pressure in the early growth stages.

### I. Relationship between the two drought tolerance traits: leaf osmotic potential at full turgor and leaf dry matter content

The *π*_*o*_ and LDMC were negatively correlated irrespective of production regime (dryland vs. irrigated) or growth stage (early vs.peak bloom (Figure 2A,B). However, the regression slope for the dryland site was significantly shallower at the early bloom stage than the peak bloom stage (*p <* 0.0001). The regression slope for the irrigated site also had significant changes during the season, but not as dramatic as in the dryland site (*p <* 0.045). As a result, the contrast in the magnitude of the regression slope between the dryland and irrigated sites became diminished at the peak bloom stage (Figure 2B). A common regression line for the peak bloom stage was found to be *π*_*o*_ = − 6.199 LDMC − 0.2285 (*R*^2^ = 0.42). The seasonal differences in regression slopes may be interpreted as being related to the osmotic and dry matter adjustments that conferred drought tolerance to the cotton plants, especially under dryland production regime.

**Fig. 2.**
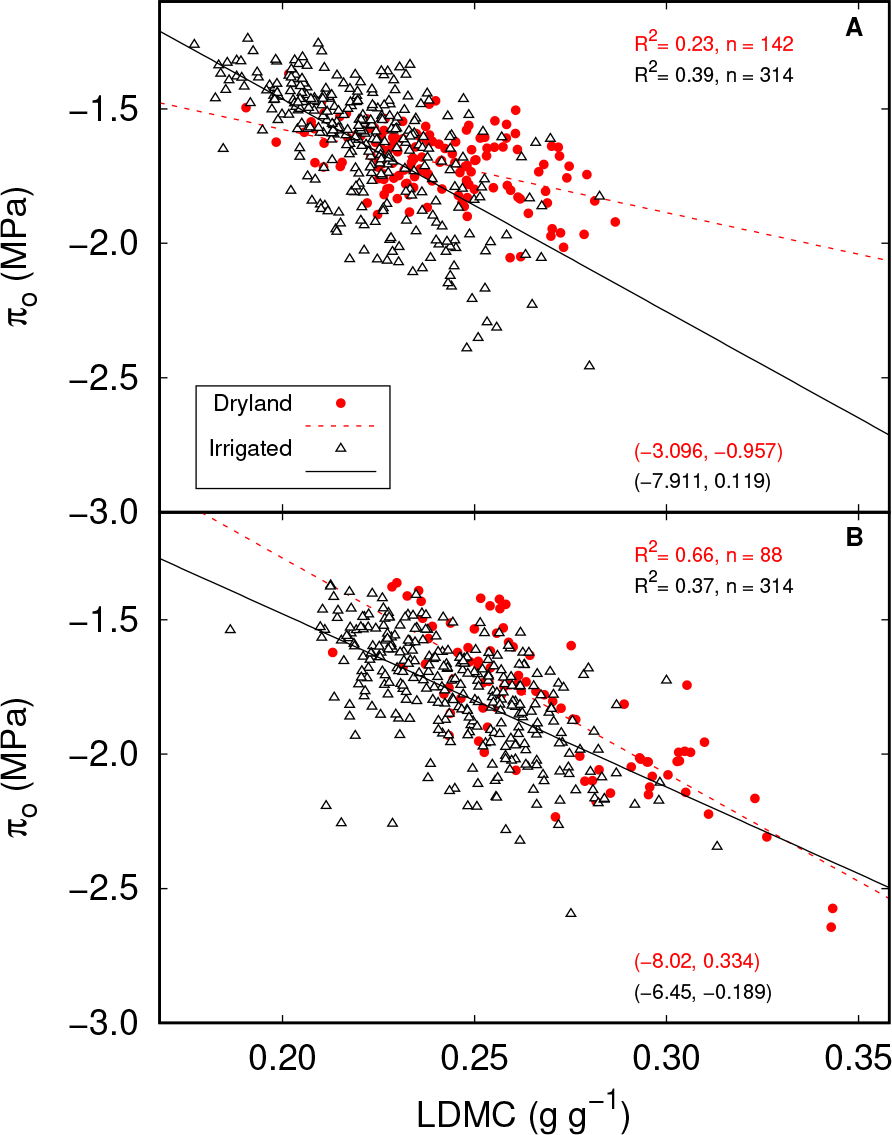
Relationships between leaf osmotic potential at full turgor (*π*_*o*_) and leaf dry matter content (LDMC) for cotton leaves measured in 2020–2022 under dryland and irrigated production conditions. The results are shown separately for the early boom stage (**A**, slopes different, *p* < 0.0001)) and peak bloom stage (**B**, slopes not different, *p* = 0.053)). Each data point represents the average of three leaves collected at one plot at a specific growth stage, site and year. The slopes and *y*-intercepts of individual regression equations are shown in parentheses in the sub-figures (slope, *y*-intercept).

### J. Responses of the two drought tolerance traits to soil water stress

The estimated water stress levels at different field sites matching the time period of our 1st and 2nd leaf sampling from 2020 to 2022 are shown in Supplementary Table A5. The Taylor site (dryland) had the highest number of days when the soil water content at one of the soil depths fell below the wilting point. However, the most severe water stress was calculated for the Uvalde Farm #2 field (SWSI = 0.608) during the 2nd sampling period in 2021. We had anticipated to see a negative correlation between SWSI and *π*_*o*_ and a positive one between SWSI and LDMC. As seen in Figure 3, however, the negative correlation between SWSI and *π*_*o*_ is not significant (*p* = 0.553), while the positive correlation between SWSI and LDMC is highly significant (*p* = 0.03). The fact that the accumulated water stress by the cotton plants during the growing season was reflected in leaf dry matter content but not in leaf osmotic potential suggests that LDMC appears to be a more sensitive drought tolerance trais than *π*_*o*_. This has some practical implications, since LDMC is much easier to measure, with a speed of measurement being *>* 10 times as fast as that of *π*_*o*_.

**Fig. 3.**
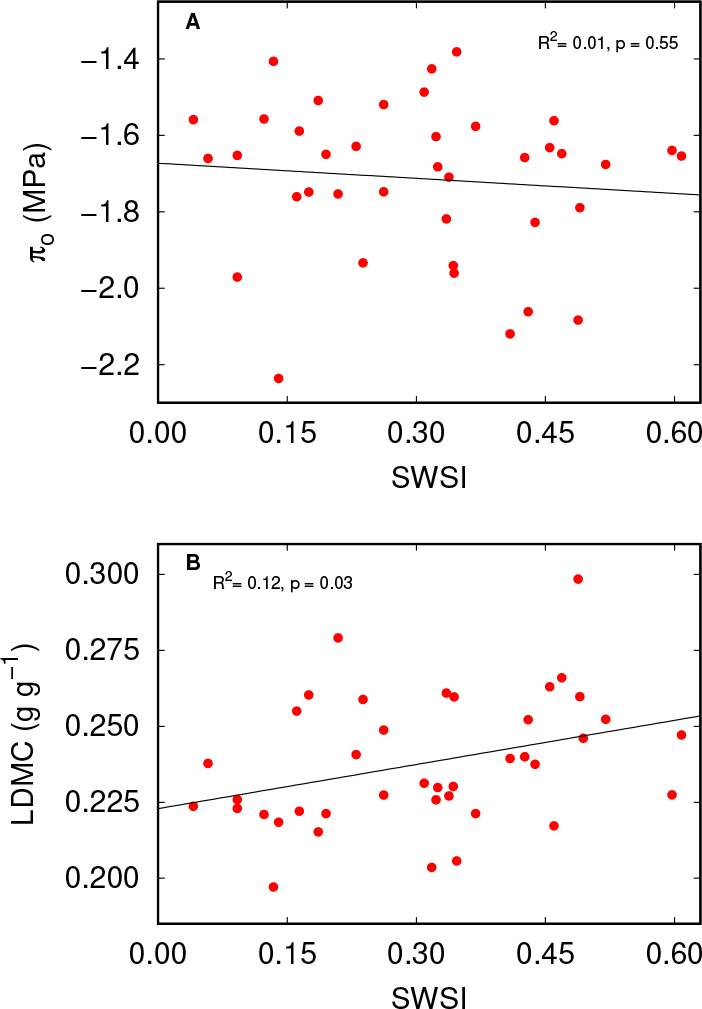
Relationships between leaf osmotic potential at full turgor (*π*_*o*_) and soil water stress index (SWSI) (**A**) and between leaf dry matter content (LDMC) and SWSI (**B**) for cotton. Each data point represents the accumulated SWSI calculated according to soil water contents measured at 15 cm, 30 cm and 60 cm soil depths at the early or peak bloom stage at one of seven sites/treatments in southwest Texas in a particular year from 2020 to 2022.

### K. Relationship between drought tolerance traits and cotton lint yield and fiber quality indices

When data at early bloom stage (same trend for the peak bloom stage data, not shown) from the dryland (Taylor) and irrigated site (Lytle) were combined, some general trends can be identified (Figures 4 and 5). A more negative *π*_*o*_ corresponded to a reduction in lint yield, as well as in micronaire, length, strength, and uniformity of fibers. But a more negative *π*_*o*_ was correlated with a high turnout percentage (Figure 4B), suggesting that increased leaf drought tolerance would possibly lead to reduced investment in cotton seeds. The linear correlations between LDMC and various yield/quality indicators were in opposite directions to those using *π*_*o*_, but in several cases, i.e., fiber uniformity, strength, length, turnout %, yielded slightly higher *R*^2^ values than with *π*_*o*_ (Figure 5).

**Fig. 4.**
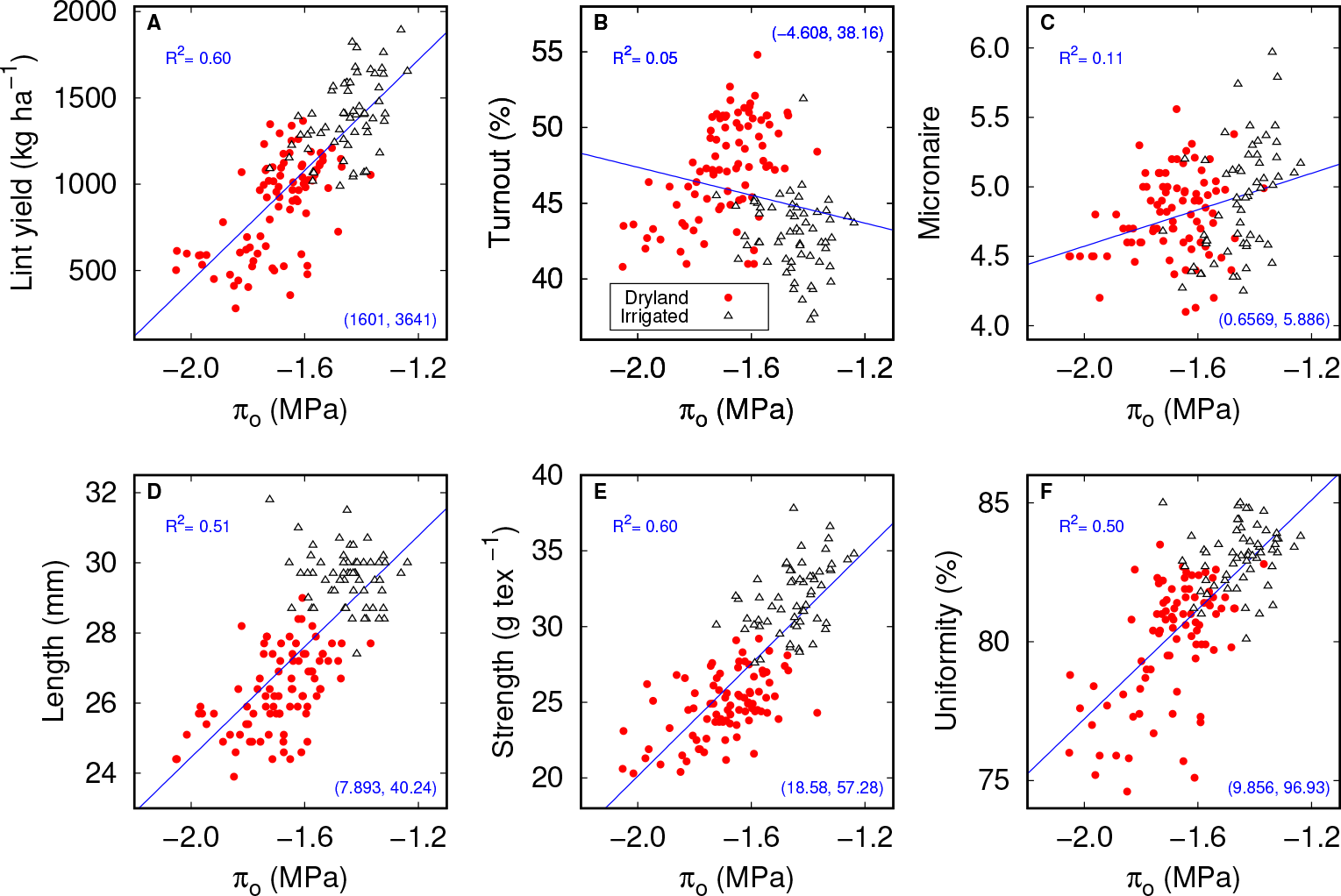
Relationships between lint yield and fiber quality indices with leaf osmotic potential at full turgor (*π*_*o*_) measured at early bloom stage in 2020–2022 under dryland and irrigated production conditions. The slope and *y*-intercept of the regression line in each sub-figure are enclosed in parentheses (slope, *y*-intercept).

**Fig. 5.**
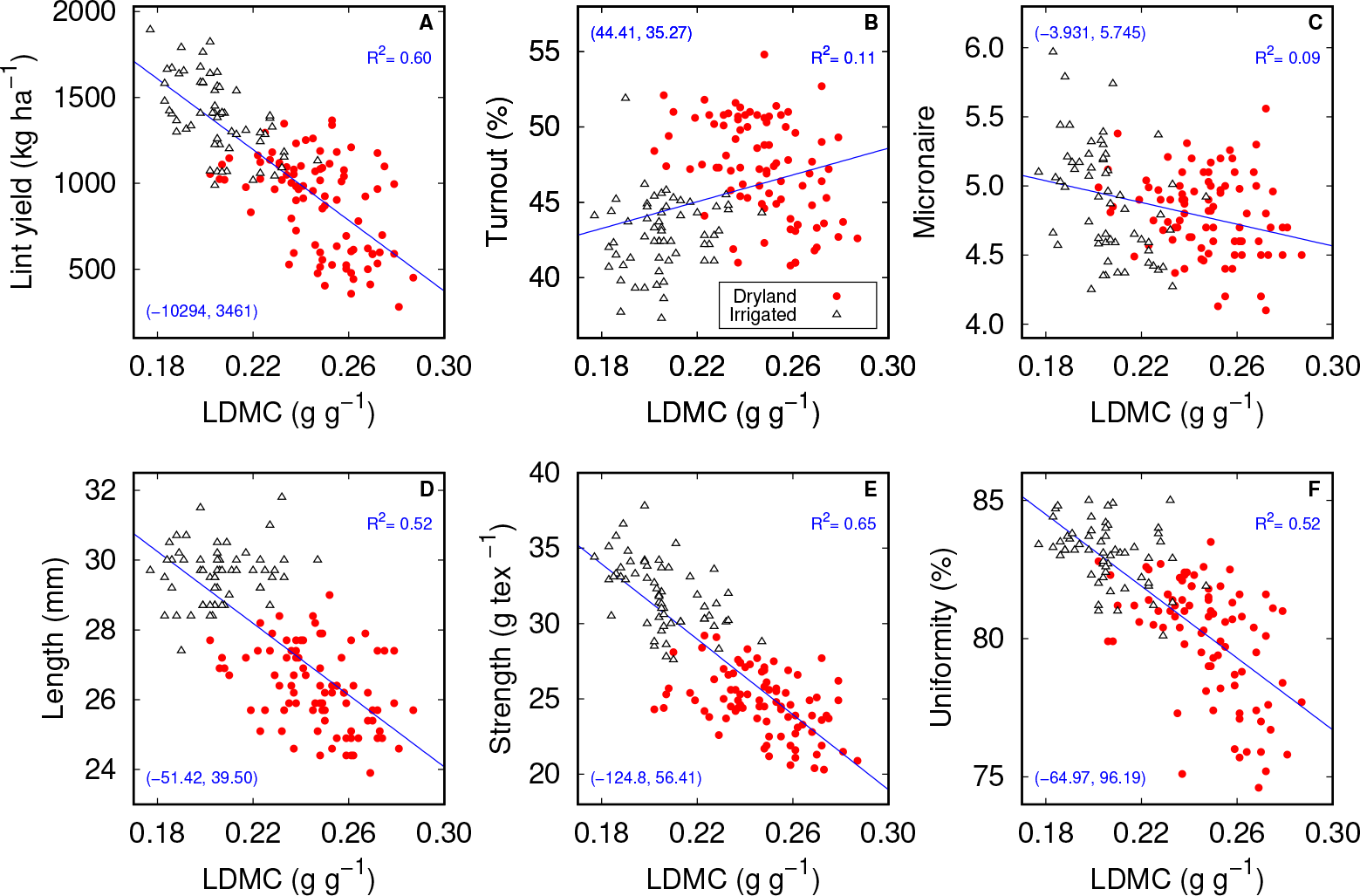
Relationships between lint yield and fiber quality indices with leaf dry matter content (LDMC) measured at early bloom stage in 2020–2022 under dryland and irrigated production conditions.The slope and *y*-intercept of the regression line in each sub-figure are enclosed in parentheses (slope, *y*-intercept).

Cotton varieties having a higher lint yield in dryland production conditions tended to not adjust their drought tolerance traits (Supplementary Figure B1). Conversely, those varieties that displayed a larger adjustment, i.e., showing a more negative *π*_*o*_ or higher LDMC under dryland vs. irrigation conditions, tended to have a lower lint yield, but potentially might exhibit a stronger capability of surviving the extreme drought.

The parameters *a*^*D*^ and *a*^*π*^ represent the adjustments in LDMC and *π*_*o*_, respectively, but the values are scaled due to the normalization of the data. As the *negative* of *π*_*o*_ was used in data normalization, the scaled values of *a*^*D*^ and *a*^*π*^ are comparable in magnitude, indicating drought tolerance capacity of the cotton varieties (the higher the number, the higher the drought tolerance capacity). It can be seen that the mean value and 95% credible intervals of *a*^*D*^ and *a*^*π*^ closely mirror those of the joint ranking results. In a further analysis as shown in Supplementary Figures C1 and C2, we can see that the 50% credible intervals of *a*^*D*^ and *a*^*π*^ for the separate analyses basically follow the trend of the joint analyses, yet the credible intervals for some varieties are wider, and especially in parameter *a*^*π*^ (such as for H 959 and NG 5150). Since the MCMC sampling procedure is based on the standardized data of LDMC and *π*_*o*_, the general consistency in drought tolerance ranking as shown in Supplementary Figures C1-C2 suggest that LDMC is equally important and effective in indicating cotton leaf drought tolerance as compared with *π*_*o*_.

The last column of Table 1 shows the lint yields of different cotton varieties measured at the dryland site. A low lint yield is generally recorded for cotton varieties with a higher drought tolerance capacity, and vice versa. Yet, an interesting exception is seen in two PhytoGen varieties, namely, PHY 332 and PHY 480, in which both the values of lint yield and drought tolerance capacity were high.

**Table 1.** Drought tolerance ranking analysis of 17 cotton varieties based on LDMC and *π*_*o*_. The 2nd to 4th columns give the posterior mean estimates of ranking, ***a***^D^ and ***a***^*π*^ for all varieties; 95% credible intervals are given in parentheses. The parameter ***a***^D^ (resp. ***a***^*π*^) is the average difference between the LDMC (resp. *π*_*o*_) measurements at dryland and irrigated sites. A larger ranking indicates better drought tolerance. Shown also are the measured lint yields (unit: kg ha^−1^) of the 17 cotton varieties under the dryland production condition.

| Variety | Ranking | $\alpha^D$ | $\alpha^\pi$ | Yield |
| --- | --- | --- | --- | --- |
| NG 4936 | 2.47 (1, 6) | 0.50 (0.18, 0.84) | 0.42 (0.05, 0.79) | 1191 |
| DP 1646 | 3.08 (1, 7) | 0.51 (0.12, 0.85) | 0.54 (0.19, 0.90) | 1178 |
| ST 5707 | 4.06 (1, 8) | 0.65 (0.20, 1.07) | 0.58 (0.13, 0.99) | 1123 |
| ST 4990 | 4.30 (1, 8) | 0.68 (0.22, 1.10) | 0.60 (0.12, 1.01) | 1019 |
| H 959 | 4.58 (1, 9) | 0.68 (0.18, 1.10) | 0.66 (0.17, 1.10) | 1381 |
| NG 4098 | 5.19 (1, 9) | 0.77 (0.33, 1.18) | 0.68 (0.18, 1.10) | 1131 |
| NG 5150 | 5.77 (1, 10) | 0.79 (0.24, 1.28) | 0.76 (0.20, 1.24) | 1425 |
| PHY 400 | 8.52 (6, 11) | 1.07 (0.81, 1.31) | 1.00 (0.72, 1.26) | 1210 |
| ST 4550 | 8.00 (4, 12) | 1.04 (0.62, 1.45) | 0.93 (0.54, 1.31) | 1086 |
| DP 2020 | 9.96 (8, 13) | 1.20 (0.93, 1.45) | 1.13 (0.87, 1.39) | 952 |
| NG 4190 | 12.25 (8, 16) | 1.43 (1.04, 1.89) | 1.33 (0.95, 1.76) | 565 |
| DG 3456 | 12.53 (9, 16) | 1.49 (1.06, 1.96) | 1.32 (0.91, 1.72) | 387 |
| PHY 332 | 12.84 (10, 16) | 1.47 (1.14, 1.82) | 1.39 (1.07, 1.78) | 1067 |
| PHY 480 | 13.88 (10, 17) | 1.60 (1.19, 2.06) | 1.48 (1.09, 1.95) | 1347 |
| DP 2012 | 14.56 (11, 17) | 1.68 (1.24, 2.14) | 1.54 (1.13, 2.00) | 550 |
| DG 3555 | 14.98 (11, 17) | 1.75 (1.29, 2.23) | 1.57 (1.16, 1.99) | 458 |
| ST 4595 | 16.05 (13, 17) | 1.87 (1.42, 2.32) | 1.75 (1.31, 2.24) | 596 |

### L. Rankings of cotton varieties for drought tolerance

The ranking results of our Bayesian hierarchical model (see Supplementary Material C for detail) are shown in Table 1. The column “Ranking” arranges the 17 varieties from less to more drought tolerant based on joint ranking using *π*_*o*_ and LDMC. NG 4936 is ranked as the lowest drought tolerant variety, while ST 4595 the highest. Largely due to the small sample size, the 95% credible intervals for the rankings are relatively wide, but still clearly separate the 17 varieties into three groups of less drought tolerant (NG 4936, DP 1646, ST 5707, and ST 4990), more drought tolerant (NG 4190, DG 3456, PHY 332, PHY 480, DP 2012, DG 3555 and ST 4595), and in between (H 959, NG 4098, NG 5150, PHY 400, ST 4550, and DP 2020). The varieties falling in the “in between” group have wider 95% credible intervals in their rankings, indicating larger uncertainty in drought tolerance compared with varieties in two other groups.

### M. Diurnal changes in leaf gas exchange, water status, canopy temperature, and stomatal behavior in cotton

*T*_canopy_, *A/E*, and *A/C*_*i*_ were significantly varied (*p <* 0.05) only with time, irrigation treatment, and variety (Figure 6). The significant effect (*p <* 0.001) of time was observed in *T*_leaf_ and *F*_0_*/F*_*m*_. Contrastingly, *F*_*v*_*/F*_*m*_ showed no significant effect for different variables and their interaction (Supplementary Table D1).

**Fig. 6.**
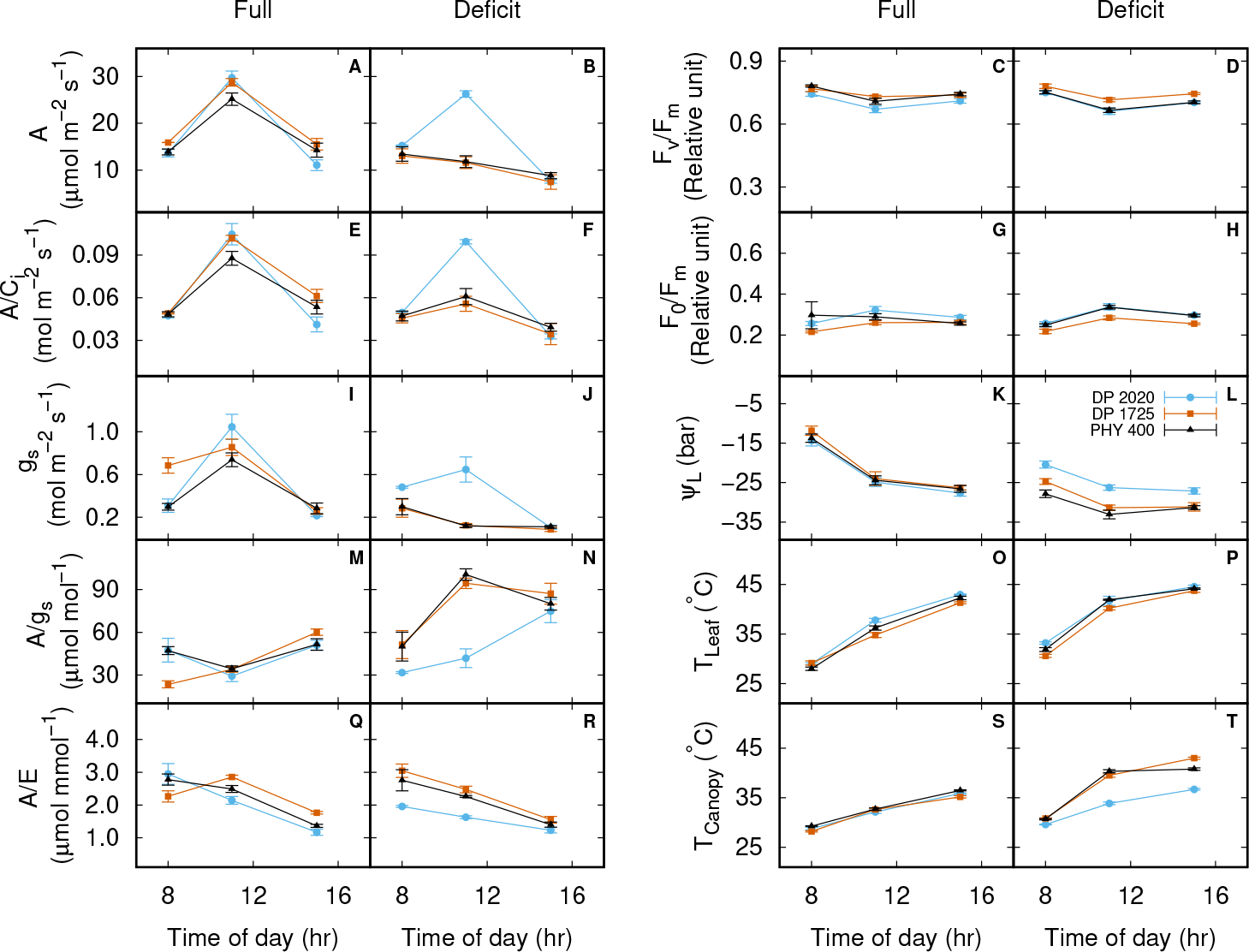
Diurnal courses of leaf physiology measurements made on three cotton genotypes, namely, DP 2020, DP 1725, and PHY 400, on July 20, 2022 at Uvalde under the full- and deficit irrigation conditions. The displayed variables include net photosynthetic rate (*A*, shown in panels **A** and **B**), carboxylation efficiency (*A/C*_*i*_, panels **E** and **J**), stomatal conductance (*g*_*s*_, panels **I** and **J**), intrinsic water use efficiency (*A/g*_*s*_, panels **M** and **N**), instantaneous water use efficiency (*A/E*, panels **U** and **R**), all measured using LI-COR 6400XT Portable Photosynthesis System on three randomly chosen sunlit mature leaves from each of the genotypes (*n* = 3, error bars indicate ± one standard errors of the means). Also shown are photochemical efficiency of PSII (*F*_*v*_*/F*_*m*_, panels **C** and **D**) and thylakoid membrane damage (*F*_0_ */F*_*m*_, panels **G** and **J**), both measured on four leaves from four different plants (*n* = 4) using OS1p Chlorophyll Fluorometer, leaf water potential (*ψL, n* = 4–6, panels **K** and **L**) measured using a pressure chamber, leaf temperature using a thermocouple with LI-6400XT Portable Photosynthesis System (*T*_leaf_, *n* = 4, panels **O** and **P**), and canopy temperature (*T*_Canopy_, *n* = 4 plants, panels **S** and **T**) measured using a hand-held infrared radiometer. The measurements were made three times at 8:00 am, 11:00 am, and 3:00 pm local time.

As expected, daytime mean *A, A/C*_*i*_, *g*_*s*_, *A/E, F*_*v*_*/F*_*m*_ and _*L*_ were reduced under water deficit (WD) treatment as compared to well-watered treatment (WW), while *A/g*_*s*_, *F*_0_*/F*_*m*_, *T*_leaf_ and *T*_canopy_ were increased. Under WW, all 3 varieties had high values of *A* and *g*_*s*_ at 11 am. However, under WD, only DP 2020 maintained high values of *A* (26.3 *μ*mol CO_2_ m^−2^ s^−1^) and *g*_*s*_ (0.647 mol H_2_O m^−2^ s^−1^), while the like values for DP 1725 and PHY 400 reduced by 55% and 81%, respectively (Figure 6B,J). These corresponded with a significantly increased *A/C*_*i*_ and reduced *A/g*_*s*_ in DP 2020 as compared with two other varieties (Figure 6F,N) at 11 am. Likewise, DP 2020 also maintained a higher *ψ*_*L*_ and cooler canopy temperature during the afternoon hours when the leaf gas exchange was measured (Figure 6L,T). The fitted parameter values of *g*_1_ of Eq. (6) show some interesting results (Table 2). In particular, the parameter *g*_1_ for DP 2020 was significantly higher than those of DP 1725 or PHY 400, which is visualized in Figure 7.

**Table 2.** Statistics of fits of the unified stomatal model (Eq. (6)) to the diurnal leaf gas exchange data of three cotton varieties measured on July 20, 2022 (a clear day) under field conditions.

| Genotype | *n* | *g*<sub>1</sub> | *s* |
| --- | --- | --- | --- |
| DP 2020 | 17 | <sup>A</sup>9.34 (0.47) | 0.099 |
| DP 1725 | 17 | <sup>BC</sup>7.72 (0.48) | 0.100 |
| PHY 400 | 18 | <sup>C</sup>7.20 (0.36) | 0.066 |

The estimated standard errors for the model parameter are shown in parentheses following the means. The unit of parameter *g*<sub>1</sub> is kPa<sup>0.5</sup>. The standard error of the regression (*s*) indicates that approximately 95% of the observations should fall within $\pm 2s$ of the predicted *g*<sub>s</sub>.
Different capital letters indicate statistically significant differences in model parameter values within the three cotton varieties (*p* < 0.05).

**Fig. 7.**
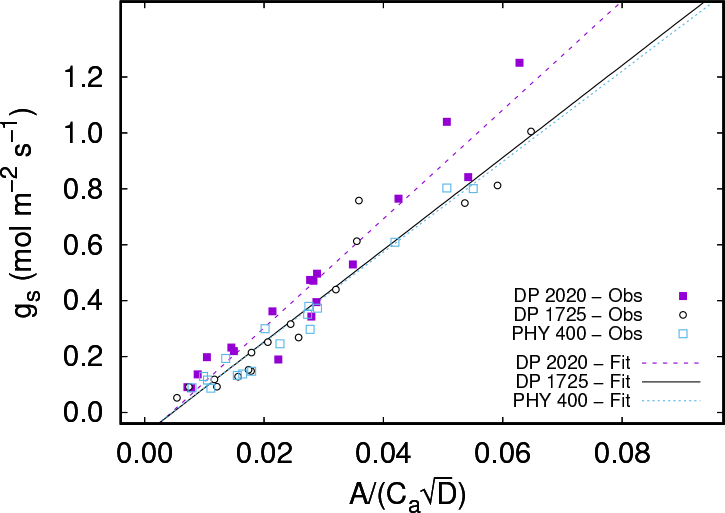
Visualization of stomatal behavior for selected three genotypes of cotton by regressing measured values of leaf stomatal conductance (*g*_*s*_, mol m^−2^ s^−1^ against 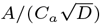. *A* is net photosynthetic rate (*μ*mol m^−2^ s^−1^), *C*_*a*_ is the atmospheric CO_2_ concentration at the leaf surface (*μ*mol mol^−1^), and *D* is leaf-to-air vapor pressure deficit (kPa). Each of the datasets of cotton included the gas exchange values of 17/18 leaves measured three times on July 20, 2022 (a clear day) at Uvalde under field condition. The slopes of the datasets are proportional to the best fits of the model (*g*_1_) as shown in Table 2. The results show that, within cotton varieties, the slope for DP 2020 was significantly higher (see Table 2 for statistical comparisons), meaning that, overall out of all leaf gas exchange data measured, DP 2020 displayed a ‘profligate’ strategy of water use, as compared with two other genotypes, in the sense of (48).

### N. Vertical root distribution and rooting depth

The differences among cotton varieties in most of the calculated root parameters are not statistically significant (*n* = 3), except in *R*_*t*2*b*_, where the estimated value for NG 4190 was significantly higher than those of three other varieties, meaning that the upper soil profile (within 40-cm depth) had 4 times the root length than the lower profile (Table 3). The maximum rooting depth for this variety was numerically shallower (i.e., 65.1 cm) than three other varieties, but the difference was not significant.

**Table 3.** Estimated values of parameters and derived variables of the model of (60) describing the vertical root distribution of selected four cotton varieties at the Taylor site (dryland production) at the early bloom stage in 2023. The analysis was based on root length density data obtained from soil samples collected from the top 100-cm soil depth on July 7, 2023.

| Variety | $K$ | $\alpha_r$ | $\beta$ | $R_{t2b}$ | $z_{95}$ (cm) |
| --- | --- | --- | --- | --- | --- |
| DP 2020 | 0.73 (0.14) | 0.023 (0.001) | 1.85 (0.09) | 2.77 (0.28) | 78.1 (1.39) |
| NG 4190 | 0.87 (0.13) | 0.034 (0.005) | 2.89 (0.45) | <sup>A</sup>3.84 (0.33) | 65.1 (4.77) |
| PHY 400 | 0.94 (0.11) | 0.026 (0.004) | 2.05 (0.27) | 2.75 (0.39) | 75.5 (3.93) |
| PHY 415 | 1.06 (0.22) | 0.028 (0.004) | 2.30 (0.34) | 2.84 (0.18) | 72.2 (4.81) |

Definition of variables: $K$ , $\alpha_r$ , and $\beta$ determine the rate of vertical reduction of root length in Eq. (7) and Eq. (8) in the main text; $R_{t2b}$ : ratio of total root length between top and bottom soil profile; $z_{95}$ : soil depth above which 95% of roots is located. The estimated standard errors for the model parameter are shown in parentheses following the means. Capital letter "A" indicates a significantly higher value as compared with the mean value for the cotton varieties ( $\alpha < 0.10$ , $n = 3$ ).

## Discussion

Leaf dry matter content (LDMC) has been used as part of a suite of leaf traits to indicate both drought tolerance (28, 33, 35, 44) and herbivore defense (24) of plants. Echoing (24)’s recent demonstration of the strong synergies between LDMC and *π*_*o*_ in semi-arid grasslands, we propose to elevate the role ‘played’ by this ordinary drought stress trait to a level as high as *π*_*o*_, the measurement of which has become a new standard for a fast estimation of plant leaf turgor loss point _TLP_ (17, 18, 20, 22–26, 44, 45, 63). This is supported by our data in that, not only did LDMC strongly correlate with *π*_*o*_ (Figure 2), similarly and even more strongly correlate with major fiber yield and quality indices of cotton (Figures 4 and 5), it also responded more sensitively to the accumulated soil water stress (Figure 3), and led to less uncertainty when used as a single basis for cotton tolerance ranking than using *π*_*o*_ (Supplementary Figures C2). A major practical advantage of LDMC over *π*_*o*_ is that it is easier and more accurate to measure, provided cautions are exercised in using consistent protocol for leaf rehydration and measurement (37, 41). The widely accepted osmometry method, however, has its own limitations (64) and source of errors (17), despite its apparent simplicity.

(17) estimated that their osmometry method involves a thirty- to fifty fold increase in measuring speed, when compared to the traditional pressure-volume method, in estimating *ψ*_TLP_. The LDMC, if used as an alternate measure of leaf drought tolerance capacity, may lead to additional tenfold increase in measuring speed, since it only involves the measurement of the turgid and dry mass of the leaf. In addition, since the whole leaf (in cotton the area of a leaf can be 100-200 cm^2^) is measured, instead of an 8-mm diameter punched leaf disc as in the osmometry method, the results should more accurately reflect the water relations properties of an average leaf cell. In some other crops with very large leaves, such as maize, the whole leaf may not be the best choice to measure LDMC, due to the significant influence of the supporting tissues in the midrib. One immediate implication of using LDMC to characterize leaf drought tolerance is the feasibility of measuring a large number of leaves in support of the remote sensing based plant phenotyping. A recent progress was demonstrated by (11) in a grassland ecosystem, although the central focus of that study was not in drought tolerance characterization.

(26) conducted a systematic ranking of drought tolerance capacity of 120 shrubs used in urban environments. To our knowledge, our work is the first effort in statistically ranking current varieties of an agriculturally important row crop for drought tolerance. The practical implication is apparent considering the fact that, in the U.S. southwest alone, the severe drought conditions in 2022 had caused 71% abandonment of planted cotton acres (65). Reliable estimation of the relative drought tolerance for major varieties that are currently being used in the region, and other similar ones worldwide, will provide readily available information for farmers to reduce loss in their planted cotton acres in the face of continued climatic drought. In addition, the quantitative data of the extent of variations in the rankings (i.e., the 95% credible intervals, seen in Table 1) provide first-hand information for cotton breeders in their search of drought tolerance genetics.

Our Bayesian hierarchical model assumes that the standardized values of LDMC and *π*_*o*_ (namely, *A*^*D*^ and *a*^*π*^) are statistically the same in indicating cotton drought tolerance capacity, and this is supported by the synergies of both the original observations of LDMC and *π*_*o*_ (Figures 4 and 5) and their adjustments (Supplementary Figure B1). The wide 95% credible intervals in Table 1, especially for the ‘in-between’ group, are mainly due to the fact that the size of the observed data is small compared with the complexity of the model. Specifically, for each numerical variable in Eq. (4) and Eq. (5), there are 17 × 2 + 12 = 46 unknown regression coefficients and 47 unknown parameters (including the error variance; see Supplementary Material C). But we only have *n* = 280 observations of LDMC and *π*_*o*_ that are applicable as inputs to this model. For this reason, the model does not consider variation of drought tolerance ranking during the growing season, which has been shown to be important in *Acer* species (45). Yet with more data becoming available, the above factor may be added to the model in the future. The Bayesian model for drought tolerance ranking as developed in this work, with its robustness in structure and flexibility in implementation, may thus be extended for use in some other applications in agricultural and related sciences where the data structure is complex, of high dimension, and unbalanced.

The observed inverse drought tolerance-lint yield/quality relationship in cotton in our study (Figures 4 and 5) is generally consistent with the prediction of LES (66), which can be seen as the results of the trade-offs between stress tolerance and growth in plants (43, 44), for which one of the leading causes could be the widely observed inverse relationship between drought tolerance capacity and photosynthetic rate (33). This line of thinking, together with the previous findings (32, 35, 52, 53), suggests the hypothesis that the more (less) drought tolerant varieties might have a shallower (deeper) rooting depth and a less (more) effective stomatal control of water loss. We will address this hypothesis momentarily with limited field data of rooting depth and leaf gas exchange, but now let’s first discuss about an anomaly in Table 1: two drought tolerant varieties, namely, PHY 332 and PHY 480, showed a high lint yield of greater than 1,000 kg ha^−1^ under dryland production conditions, a trend contradicting the prediction of LES. We don’t have an explanation for the ‘behavior’ of these two varieties, but they may possess some unique ways for effective water use (67), while having a high drought tolerance capacity. This needs attention for future studies.

Although *z*_95_ of NG 4190, a variety considered as drought tolerant in Table 1, was not significantly lower than DP 2020 or PHY 400 (falling in the ‘in-between’ group in Table 1), it had a significantly higher value of parameter *R*_*t*2*b*_, or a predominantly shallower root distribution, which is consistent with the prediction of LES. The roots obtained from field soil sampling might originate from both the current year and previous year’s growths. However, since the estimated values of *β* parameter of Eq. (8) are below 3.0 (Table 3), the impact from previous years’ roots is arguably limited (68, 69). But this needs to be clarified further in the future.

The eco-physiological significance of parameter *g*_1_ in Eq. (6) has been well-documented (48, 70): it is generally inversely proportional to leaf WUE. The estimated values for our cotton plants are close to the typical ones for tropical vegetation (48). Both the diurnal leaf gas exchange data and those of *g*_1_ indicate that, compared with PHY 400, DP2020 uses water in a more ‘‘profligate’ way under water deficit conditions (Figures 6 and 7). However, this does not seem to receive strong support from our drought tolerance ranking results (Table 1) and rooting depth survey (Table 3), although parameter *z*_95_ for DP 2020 was numerically deeper when compared PHY 400 and DP 1725. Future studies are needed to use additionally measured drought avoidance traits, such as rooting depth distribution and leaf gas exchange rates, to help interpret the measured leaf drought tolerance traits and crop yield performance.

Overall, the prominent role played by LDMC in cotton drought tolerance and the quantitative drought tolerance rankings of the selected cotton varieties based on our field experiments and statistical modeling may help build a foundation for future work to elucidate the eco-physiological mechanisms underlying drought stress resistance in cotton, and perhaps in other crop species as well, by targeting representative genotypes with a contrasting drought tolerance capacity.

## ACKNOWLEDGEMENTS

We thank Shane Sieckenius for drawing Supplementary Figure A1, Shane Sieckenius and Dean Hillis for crop management, Nathan Alonzo, Luke Alejandro, Mark Hernandez, Muhammad Saleem, Joe Moreno, and Steven Ramos for assistance in field and lab measurements. We also thank the following regional farmers for allowing us to use their lands for conducting cotton experiments: David Kriewald, Gary Pastushok, Justin Speer, Manuel Ortez, Nathan Philips, and Rick Kruger. The work received funding support from USDA NIFA Hatch project TEX0-1-9574, Cotton Incorporated project 20-557TX, and NSF DMS-2245591.

## Supporting Information

- **Supplementary Material A:** Study locations, summary of amounts of water input, lists of cotton variety names, and values of soil water stress index.
  – Figure A1. Locations of the six field sites in southern Texas …
  – Table A1. List of varieties in 2020.
  – Table A2. List of varieties in 2021.
  – Table A3. List of varieties in 2022.
  – Table A4. Summary of the total amounts of rain and irrigation received
  – Table A5. Values of the soil water stress index (SWSI) calculated for different sampling periods …
- **Supplementary Material B:** Relationship between dryland cotton yield and adjustments in *π*_*o*_ and LDMC.
- **Supplementary Material C:** Drought Tolerance Ranking of Cotton Varieties
  – C1: Data for drought tolerance ranking
    * Table C1: Number of replicates for LDMC measurement by cotton variety and season…
  – C2: Bayesian joint modeling of leaf dry matter content and leaf osmotic potential
  – C3: Conditional posterior distributions
  – C4: Markov chain Monte Carlo sampling
  – C5: Results
    * Figure C1: Drought tolerance ranking of 17 cotton varieties via joint Bayesian analysis of LDMC and *π*_*o*_.
    * Figure C2: Estimates of *a*^*D*^ and *a*^*π*^ from the MCMC output.
- **Supplementary Material D.** Table D1. Probability values of the ANOVA results for variables of leaf gas exchange and water relations measurements …

## Bibliography

1. P. S. Tang and J. S. Wang. A thermodynamic formulation of the water relations in an isolated living cell. Journal of Physical Chemistry, 45:443–453, 1941.

2. P. F. Scholander, H. T. Hammel, E. D. Bradstreet, and E. A. Hemmingsen. Sap pressure in vascular plants. Science, 148:339–346, 1965.

3. J. R. Philip. Plant water relations: some physical aspects. Annual Review of Plant Physiology, 17:245–268, 1966.

4. P. G. Jarvis. Water transfer in plants. In D. A. De Vries and N. H. Afgan, editors, Heat and Mass Transfer in the Biosphere, volume 1, pages 369–394, Washington, D. C., 1975. Scripta Book, Co.

5. R. Oren, J. S. Sperry, G. G. Katul, D. E. Pataki, B. E. Ewers, N. Phillips, and K. V. R. Schäfer. Survey and synthesis of intra- and interspecific variations in stomatal sensitivity to vapor pressure deficit. Plant, Cell and Environment, 22:1515–1526, 1999.

6. F. Tardieu and T. Simonneau. Variability among species of stomatal control under fluctuating soil water status and evaporative demand: modeling isohydric and anisohydric behaviours. Journal of Experimental Botany, 49:419–432, 1998.

7. J. M. Norman and M. C. Anderson. Soil-plant-atmosphere continuum. In D. Hillel, editor, Encyclopedia of Soils in the Environment, pages 513–521. Elsevier, Oxford, 2005.

8. J. Landsberg and R. Waring. Water relations in tree physiology: where to from here? Tree Physiology, 37:18–32, 2016.

9. M. S. Moran, T. R. Clarke, Y. Inoue, and A. Vidale. Estimating crop water deficit using the relation between surface–air temperature and spectral vegetation index. Remote Sensing of Environment, 49:246–263, 1994.

10. S. Elsayed, B. Mistele, and U. Schmidhalter. Can changes in leaf water potential be assessed spectrally? Functional Plant Biology, 38:523–533, 2011.

11. H. W. Polley, C. Yang, B. J. Wilsey, and P. A. Fay. Spectrally derived values of community leaf dry matter content link shifts in grassland composition with changes in biomass production. Remote Sensing in Ecolology and Conservation, 6:344–353, 2020.

12. X. Dong, B. Peng, S. Sieckenius, R. Raman, M. M. Conley, and D. I. Leskovar. Leaf water potential of field crops estimated using NDVI in ground-based remote sensing - opportunities to increase prediction precision. PeerJ, 9:e12005, 2021.

13. M. Yoosefzadeh-Najafabadi, K. D. Singh, A. Pourreza, K. S. Sandhu, A. Adak, S. C. Murray, M. Eskandari, and I. Rajcan. Remote and proximal sensing: How far has it come to help plant breeders? Advances in Agronomy, page 10.1016/bs.agron.2023.05.004, 2023.

14. T. M. Hinckley, F. Duhme, A. R. Hinckley, and H. Richter. Water relations of drought hardy shrubs: Osmotic potential and stomatal reactivity. Plant, Cell and Environment, 3:131–140, 1980.

15. X. Dong and X. Zhang. Some observations of the adaptations of sandy shrubs to the arid environment in the Mu Us Sandland: Leaf water relations and anatomic features. Journal of Arid Environments, 48:41–48, 2001.

16. Frederick C. Meinzer, David R. Woodruff, Danielle E. Marias, Katherine A. McCulloh, and Sanna Sevanto. Dynamics of leaf water relations components in co-occurring iso- and anisohydric conifer species. Plant, Cell and Environment, 37:2577–2586, 2014.

17. Megan K. Bartlett, Ya Zhang, Christine Scoffoni, Shanwen Sun, Rico Ardy, Kunfang Cao, and Lawren Sack. Rapid determination of comparative drought tolerance traits: using an osmometer to predict turgor loss point. Methods in Ecology and Evolution, 3:880–888, 2012.

18. I. Maréchaux, M. K. Bartlett, L. Sack, C. Baraloto, J. Engel, E. Joetzjer, and J. Chave. Drought tolerance as predicted by leaf water potential at turgor loss point varies strongly across species within an Amazonian forest. Functional Ecology, 29:1268–1277, 2015.

19. H. Sjöman, A. D. Hirons, and N. L. Bassuk. Urban forest resilience through tree selection — Variation in drought tolerance in Acer. Urban Forestry & Urban Greening, 14:858–865, 2015.

20. K. B. Mart, E. J. Veneklaas, and H. Bramley. Osmotic potential at full turgor: an easily measurable trait to help breeders select for drought tolerance in wheat. Plant Breeding, 135:279–285, 2016.

21. J. M. Banks and A. D. Hirons. Alternative methods of estimating the water potential at turgor loss point in Acer genotypes. Plant Methods, 15:34, 2019.

22. Robert J. Griffin-Nolan, Troy W. Ocheltree, Kevin E. Mueller, Dana M. Blumenthal, Julie A. Kray, and Alan K. Knapp. Extending the osmometer method for assessing drought tolerance in herbaceous species. Oecologia, 189:353–363, 2019.

23. F. Petruzzellis, T. Savi, G. Bacaro, and A. Nardini. A simplified framework for fast and reliable measurements of leaf turgor loss point. Plant Physiology and Biochemistry, 139:395–399, 2019.

24. D. Blumenthal, K. E. Mueller, J. A. Kray, T. W. Ocheltree, D. J. Augustine, and K. R. Wilcox. Traits link drought resistance with herbivore defence and plant economics in semi-arid grasslands: The central roles of phenology and leaf dry matter content. Journal of Ecology, 108:2336–2351, 2020.

25. D. M. Blumenthal, D. R. LeCain, L. M. Porensky, E. A. Leger, R. Gaffney, T. W. Ocheltree, and A. M. Pilmanis. Local adaptation to precipitation in the perennial grass Elymus elymoides: Trade-offs between growth and drought tolerance traits. Evolutionary Applications, 14:524–535, 2021.

26. H. Sjöman, S. Ignell, and A. Hirons. Selection of shrubs for urban environments — An evaluation of drought tolerance of 120 species and cultivars. HortScience, 58:573–579, 2023.

27. Y. N. S. Cheung, M. T. Tyree, and J. Dainty. Water relations parameters on single leaves obtained in a pressure bomb and some ecological interpretations. Canadian Journal of Botany, 53:1342–1346, 1975.

28. S. C. Clifford, S. K. Arndt, J. E. Corlett, S. Joshi, N. Sankhla, M. Popp, and H. G. Jones. The role of solute accumulation, osmotic adjustment and changes in cell wall elasticity in drought tolerance in Ziziphus mauritiana (Lamk.). Journal of Experimental Botany, 49:967–977, 1998.

29. Q. Gao, P. Zhao, X. Zeng, X. Cai, and W. Shen. A model of stomatal conductance to quantify the relationship between leaf transpiration, microclimate and soil water stress. Plant, Cell and Environment, 25:1373–1381, 2002.

30. T. I. Lenz, I. J. Wright, and M. Westoby. Interrelations among pressure-volume curve traits across species and water availability gradients. Physiologia Plantarum, 127:423–433, 2006.

31. Megan K. Bartlett, Christine Scoffoni, and Lawren Sack. The determinants of leaf turgor loss point and prediction of drought tolerance of species and biomes: a global meta-analysis. Ecology Letters, 15:393–405, 2012.

32. S. Fan, T. J. Blake, and E. Blumwald. The relative contribution of elastic and osmotic adjustments to turgor maintenance of woody species. Physiologia Plantarum, 90:408–413, 1994.

33. Ü. Niinemets. Global-scale climatic controls of leaf dry mass per area, density, and thickness in trees and shrubs. Ecology, 82:453–469, 2001.

34. R. K. Monson and S. D. Smith. Seasonal water potential components of Sonoran Desert plants. Ecology, 63:113–123, 1982.

35. S. D. Davis and H. A. Mooney. Tissue water relations for four co-occurring chaparral shrubs. Oecologia, 70:527–535, 1986.

36. K. J. Niklas. Mechanical behavior of plant tissues as inferred from the theory of perssureized celullar solids. American Journal of Botany, 76:929–937, 1989.

37. E. Garnier, B. Shipley, C. Roumet, and G. Laurent. A standardized protocol for the determination of specific leaf area and leaf dry matter content. Functional Ecology, 15:688–695, 2001.

38. X. Dong, B. Patton, P. Nyren, R. Limb, L. Cihacek, D. Kirby, and E. Deckard. Leaf-water relations of a native and an introduced grass species in the mixed-grass prairie under cattle grazing. Applied Ecology and Environmental Research, 9:311–331, 2011.

39. E. Garnier and G. Laurent. Leaf anatomy, specific mass and water content in congeneric annual and perennial grass species. New Phytologist, 128:725–736, 1994.

40. Ü. Niinemets. Research review: Components of leaf dry mass per area – thickness and density – alter leaf photosynthetic capacity in reverse directions in woody plants. New Phytologist, 144:35–47, 1999.

41. Bill Shipley and Thi-Tam Vu. Dry matter content as a measure of dry matter concentration in plants and their parts. New Phytologist, 153:359–364, 2002.

42. I. J. Wright, P. B. Reich, M. Westoby, and et al. The worldwide leaf economics spectrum. Nature, 428:821–827, 2004.

43. P. B. Reich. The worldwide ’fast-slow’ plant economics spectrum: a traits manifesto. Journal of Ecology, 102:275–301, 2014.

44. Shi-Dan Zhu, Ya-Jun Chen, Qing Ye, Peng-Cheng He, Hui Liu, Rong-Hua Li, Pei-Li Fu, Guo-Feng Jiang, and Kun-Fang Cao. Leaf turgor loss point is correlated with drought tolerance and leaf carbon economics traits. Tree Physiology, 38:658–663, 2018.

45. J. M. Banks, G. C. Percival, and G. Rose. Variations in seasonal drought tolerance rankings. Trees: Structure and Function, 33:1063–1072, 2019.

46. E.-D. Schulze, R. H. Robichaux, J. Grace, P. W. Rundel, and J. R. Ehleringer. Plant water balance. Bioscience, 37:30–37, 1987.

47. J. T. Ball, I. E. Woodrow, and J. A. Berry. A model predicting stomatal conductance and its contribution to the control of photosynthesis under different environmental conditions. In J. Biggens, editor, Progress in Photosynthesis Research: Proceedings of the VIIth International Congress on Photosynthesis, volume 4, pages 221–224, Dordrecht, The Netherlands, 1987. Martinus Nijhoff Publishers.

48. B. E. Medlyn, R. A. Duursma, D. Eamus, D. S. Ellsworth, I. C. Prentice, C. V. M. Barton, K. Y. Crous, P. De Angelis, M. Freeman, and L. Wingate. Reconciling the optimal and empirical approaches to modelling stomatal conductance. Global Change Biology, 17:2134–2144, 2011.

49. P. G. Jarvis. The interpretation of the variations in leaf water potential and stomatal conductance found in canopies in the field. Philosophical Transactions of the Royal Society of London. Series B: Biological, 273:593–610, 1976.

50. I. R. Cowan and G. D. Farquhar. Stomatal function in relation to leaf metabolism and environment. In Integration of activity in the higher plant, Soc. Exp. Biol. Symp., number 31 in XXXI, pages 471–505. Cambridge University Press, New York, 1977.

51. J. Canadell, R. B. Jackson, J. R. Ehleringer, H. A. Mooney, O. E. Sala, and E.-D. Schulze. Maximum rooting depth of vegetation types at the global scale. Oecologia, 108:583–595, 1996.

52. D. K. Poole and P. C. Miller. Water relations of selected species of chaparral and coastal sage communities. Ecology, 56:1118–1128, 1975.

53. D. S. Crombie, J. T. Tippett, and T. C. Hill. Dawn water potential and root depth of trees and understory species in Southwestern Australia. Australian Journal of Botany, 36:621–631, 1988.

54. M. M. Thornton, R. Shrestha, Y. Wei, P. E. Thornton, S. Kao, and B. E. Wilson. Daymet: Daily surface weather data on a 1-km grid for North America. Version 4. ORNL DAAC, Oak Ridge, Tennessee, USA. 10.3334/ORNLDAAC/1840, 2020.

55. H. Rymbai, R. H. Laxman, M. R. Dinesh, V. S. J. Sunoj, K. V. Ravishankar, and A. K. Jha. Diversity in leaf morphology and physiological characteristics among mango (Mangifera indica) cultivars popular in different agro-climatic regions of India. Scientia Horticulturae, 176: 189–193, 2014.

56. D. M. Oosterhuis. Growth and development of a cotton plant. In W. N. Miley and D. M. Oosterhuis, editors, Nitrogen Nutrition in Cotton: Practical Issues, Proceedings of Southern Branch Workshop for Practicing Agronomists, pages 1–24, Madison, WI, 1990. American Society of Agronomy.

57. Pertti Hari, Annikki Mäkelä, Eeva Korpilahti, and Maria Holmberg. Optimal control of gas exchange. Tree Physiology, 2:169–175, 1986.

58. Remko A. Duursma, Christopher J. Blackman, Rosana Lopéz, Nicolas K. Martin-StPaul, Hervé Cochard, and Belinda E. Medlyn. On the minimum leaf conductance: its role in models of plant water use, and ecological and environmental controls. New Phytologist, 221:693–705, 2019.

59. A. Gardner, M. Jiang, D. S. Ellsworth, A. R. MacKenzie, J. Pritchard, M. K.-F. Bader, C. V. M. Barton, C. Bernacchi, C. Calfapietra, K. Y. Crous, M. E. Dusenge, T. E. Gimeno, M. Hall, S. Lamba, S. Leuzinger, J. Uddling, J. Warren, G. Wallin, and B. E. Medlyn. Optimal stomatal theory predicts CO2 responses of stomatal conductance in both gymnosperm and angiosperm trees. New Phytologist, 237:1229–1241, 2023.

60. X. Dong, B. Peng, X. Liu, K. Qin, Q. Xue, and D. I. Leskovar. An automated calculation of plant root distribution parameters based on root length density data. Applied Ecology and Environmental Research, 17:3545–3552, 2019.

61. J. Wu, R. Zhang, and S. Gui. Modeling soil water movement with water uptake by roots. Plant and Soil, 215:7–17, 1999.

62. C. S. P. Ojha and A. K. Rai. Nonlinear root water uptake model. Journal of Irrigation and Drainage Engineering, 122:198–201, 1996.

63. M. Májeková, J. Martínková, and T. Hájek. Grassland plants show no relationship between leaf drought tolerance and soil moisture affinity, but rapidly adjust to changes in soil moisture. Functional Ecology, 33:774–785, 2019.

64. T. E. Sweeney and C. A. Beuchat. Limitations of methods of osmometry: measuring the osmolality of biological fluids. American Journal of Physiology, 264:R469–R480, 1993.

65. L. Meyer, T. Dew, M. Grace, K. Lanclos, S. MacDonald, and G. Soley. The world and United States cotton outlook. https://www.usda.gov/sites/default/files/documents/2023AOF-cotton-outlook.pdf, 2023.

66. I. J. Wright, P. B. Reich, M. Westoby, D. D. Ackerly, Zdravko Baruch, Frans Bongers, Jeannine Cavender-Bares, Terry Chapin, Johannes H. C. Cornelissen, Matthias Diemer, Jaume Flexas, Eric Garnier, Philip K. Groom, Javier Gulias, Kouki Hikosaka, Byron B. Lamont, Tali Lee, William Lee, Christopher Lusk, Jeremy J. Midgley, M.-L. Navas, Ü. Niinemets, Jacek Oleksyn, Noriyuki Osada, Hendrik Poorter, Pieter Poot, Lynda Prior, Vladimir I. Pyankov, Catherine Roumet, Sean C. Thomas, Mark G. Tjoelker, Erik J. Veneklaas, and Rafael Villar. The worldwide leaf economics spectrum. Nature, 428:821–827, 2004.

67. A. Blum. Effective use of water (EUW) and not water-use efficiency (WUE) is the target of crop yield improvement under drought stress. Field Crops Reseach, 112:119–123, 2009.

68. R. J. Lorenz and G. A. Rogler. Grazing and fertilization affect root development of range grasses. Journal of Range Management, 20:129–132, 1967.

69. X. Dong, B. D. Patton, A. C. Nyren, P. E. Nyren, and L. D. Prunty. Quantifying root water extraction by rangeland plants through soil water modeling. Plant and Soil, 335:181–198, 2010.

70. A. Stefanski, E. E. Butler, R. Bermudez, R. A. Montgomery, and P. B. Reich. Stomatal behaviour moderates the water cost of CO2 acquisition for 21 boreal and temperate species under experimental climate change. Plant, Cell and Environment, page 1–18. 10.1111/pce.14559, 2023.

