## Supplementary Material for "Rapid measurement and statistical ranking of leaf drought tolerance capacity in cotton"

**Supplementary Material A.** Study locations, summary of amounts of water input, lists of cotton variety names, and values of soil water stress index.

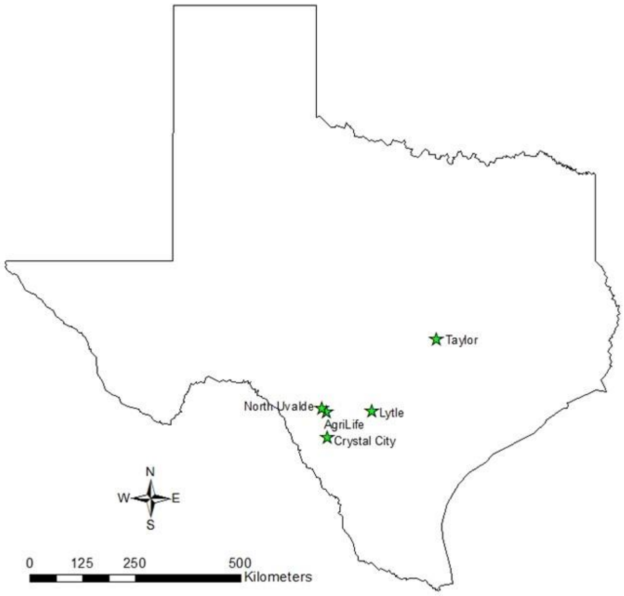


Figure A1. Locations of the six field sites in southern Texas that were used for cotton leaf sampling from 2020 to 2022: one site in Crystal City (average annual precipitation 523 mm), three sites in Uvalde (594 mm), one in Lytle (683 mm), and one in Taylor (892 mm).

Table A1. List of varieties in 2020.

| ID | Variety | Site | ID | Variety | Site | ID | Variety | Site |
|---|---|---|---|---|---|---|---|---|
| 1 | 19R132 B3XF | T | 16 | DG 3421 B3XF | UR,T | 31 | NG 4936 B3XF | L, UR, T |
| 2 | 19R237 B3XF | T | 17 | DG 3615 B3XF | L, UR | 32 | NG 5007 B2XF | UR |
| 3 | 20R 734 B3XF | T | 18 | DP 1044 B2RF | UR | 33 | NG 5711 B3XF | U2 |
| 4 | 20R 741 B3XF | T | 19 | DP 1646 B2XF | L, UR, T | 34 | PHY 340 W3FE | UR |
| 5 | 20R 743 B3XF | T | 20 | DP 1725 B2XF | UR | 35 | PHY 400 W3FE | L, T |
| 6 | 20R 749 B3XF | T | 21 | DP 1820 B3XF | T | 36 | PHY 480 W3FE | L, UR |
| 7 | 20R 750 B3XF | T | 22 | DP 1845 B3XF | T | 37 | ST 4550 GLTP | L, T |
| 8 | 20R 752 B3XF | T | 23 | DP 1865 B3XF | C, U1 | 38 | ST 4848 GLT | UR |
| 9 | BX 2116 GLTP | T | 24 | DP 1948 B3XF | T | 39 | ST 4949 GLT | UR |
| 10 | BX 2141 GLTP | T | 25 | DP 2020 B3XF | L, T | 40 | ST 4990 B3XF | L, UR, T |
| 11 | BX 2191 B3XF | T | 26 | DP 2044 B3XF | T | 41 | ST 5600 NR B2XF | T |
| 12 | BX 2192 B3XF | T | 27 | FM 1953 GLTP | UR | 42 | ST 5610 B3XF | T |
| 13 | BX 2193 B3XF | T | 28 | FM 2398 GLTP | T | 43 | ST 5707 B2XF | L, T |
| 14 | BX 2194 B3XF | T | 29 | FM 4480 B3XF | T | | | |
| 15 | CG 3885 B2XF | UR | 30 | NG 4098 B3XF | L, UR, T | | | |

C-Crystal City; L-Lytle; T-Taylor; UR-Uvalde Research; U1-Uvalde farm #1 (UJ); U2-Uvalde farm #2 (UN)

Table A2. List of varieties in 2021.

| ID | Variety | Site | ID | Variety | Site | ID | Variety | Site |
|---|---|---|---|---|---|---|---|---|
| 1 | DP 2012 B3XF | T | 16 | DG 3421 B3XF | UR | 31 | NG 4936 B3XF | L, UR, T |
| 2 | 20R732 B3XF | T | 17 | DG 3615 B3XF | L, UR | 32 | NG 5007 B2XF | UR |
| 3 | 20R 734 B3XF | T | 18 | DG 3402 B3XF | T | 33 | NG 5711 B3XF | U2 |
| 4 | 21R 739 B3XF | T | 19 | DP 1646 B2XF | L, UR, T | 34 | PHY 332 W3FE | L, UR, T |
| 5 | 21R 626 B3XF | T | 20 | DP 1725 B2XF | UR | 35 | PHY 400 W3FE | L, T |
| 6 | 21R 632 B3XF | T | 21 | H 959 B3XF | L, T | 36 | PHY 480 W3FE | UR |
| 7 | DP 1851 B3XF | T | 22 | DP 1845 B3XF | T | 37 | ST 4550 GLTP | T |
| 8 | DP 1845 B3XF | T | 23 | DP 1865 B3XF | C, U1 | 38 | ST 4848 GLT | UR |
| 9 | NG 5150 B3XF | L | 24 | NG 5150 B3XF | L, T | 39 | ST 4949 GLT | UR |
| 10 | PHY340W3FE | UR | 25 | DP 2020 B3XF | L, T, UR | 40 | ST 4990 B3XF | UR, T |
| 11 | BX 2295 B3XF | T | 26 | FM 1730 GLTP | T | 41 | ST 5091 B3XF | T |
| 12 | BX 2296 B3XF | T | 27 | FM 1953 GLTP | UR | 42 | ST 4993 B3XF | T |
| 13 | BX 2297 B3XF | T | 28 | FM 2398 GLTP | T | 43 | ST 5707 B2XF | T |
| 14 | BX 2298 B3XF | T | 29 | FM 2498 GLTP | T | | | |
| 15 | CG 3885 B2XF | UR | 30 | NG 4098 B3XF | UR | | | |

Table A3. List of varieties in 2022.

| ID | Variety | Site | ID | Variety | Site | ID | Variety | Site |
|---|---|---|---|---|---|---|---|---|
| 1 | DP 2012 B3XF | L,T | 10 | DG 3555 B3XF | T | 19 | PHY 332 W3FE | L, UR, T |
| 2 | NG 5007 B3XF | UR | 11 | DP 1646 B2XF | UR | 20 | PHY 400 W3FE | L, UR, T |
| 3 | PHY340 W3FE | UR | 12 | DP 1725 B2XF | UR | 21 | PHY 480 W3FE | UR |
| 4 | PHY 411 B3XF | L | 13 | DP 2015 B2XF | U1 | 22 | ST 4595 GLTP | T |
| 5 | DG 3615 B3XF | UR | 14 | DP 2020 B3XF | L,T,UR | 23 | ST 4990 B3XF | UR, T |
| 6 | DG 4098 B3XF | UR | 15 | NG 5711 GLTP | U2 | 24 | ST 5091 B3XF | L |
| 7 | DG 3456 B2XF | T | 16 | NG 4936 B3XF | UR | 25 | ST 4993 B3XF | T |
| 8 | DG 3421 B3XF | UR | 17 | NG 5007 B2XF | UR | 26 | ST 5471 B2XF | UR |
| 9 | DG 3615 B3XF | UR | 18 | NG 4190 B3XF | L, T | | | |

Table A4. Summary of the total amounts of rain and irrigation received at each of the six sites from March 13 to October 8 in 2020, 2021, 2022, as well as the average rain received during the same time period in past 43 years from 1980 to 2022, with standard deviations (sdtev) in parentheses.

| Location | *2020 (mm) | 2021 (m) | 2022 (mm) | 43-yr average/stdev (mm) |
|---|---|---|---|---|
| Taylor | †491/0/491 | 651/0/651 | 285/0/285‡ | 540 (154) |
| Lytle | 348/252/600 | 646/89/735‡ | 204/340/544 | 493 (211) |
| Uvalde Farm #1 | 312/144/456 | 472/206/678‡ | 359/221/580‡ | 389 (155) |
| Uvalde Farm #2 | 314/130/444 | 477/146/623‡ | 361/153/514 | 396 (164) |
| Crystal City | 241/54/295 | 530/51/581‡ | 366/89/455 | 337 (139) |
| Uvalde Research | 249/292/541 | 273/210/483 | 317/348/665‡ | 410 (168) |

*Total amounts of seasonal rainfall received were obtained from the 1-km grid weather data product available at https://daymet.ornl.gov (Thornton et al., 2020), and the amounts of irrigation applied at different field sites were measured using 2 standard rain gauges.

† Data are shown in the format of "rain (mm)/irrigation (mm)/total (mm)".

‡ Case where the total amount of water inputs (rain + irrigation) was either greater or less than the long-term average rainfall (± 1 stdev) received at a site.

Table A5. Values of the soil water stress index (SWSI) calculated for different sampling periods for different fields/environments in 2020, 2021 and 2022.The values lie within the range of 0 and 1, indicating minimum and maximum water stress, respectively, in terms of the relative degree of water availability to plants. The 1st sampling was done in early-peak flowering and the 2nd one was done in peak to late flowering.

| Sites/environments | 2020 1st | 2020 2nd | 2021 1st | 2021 2nd | 2022 1st | 2022 2nd |
|---|---|---|---|---|---|---|
| Crystal City (CJ) | 0.092 | 0.209 | 0.455 | 0.469 | - | - |
| Uvalde 1 (UJ) | 0.460 | 0.490 | 0.309 | 0.325(1) | 0.092 | 0.140 (18) |
| Uvalde 2 (UN) | 0.318 | 0.426 | 0.597 | 0.608 | 0.347 | 0.369 |
| Uvalde (Deficit) | 0.195 | 0.344 | 0.058 | 0.175 | 0.409 | 0.430 |
| Uvalde (Full) | 0.123 | 0.238 | 0.041 | 0.161 | 0.343 | 0.438 |
| Lytle | 0.134 | 0.323 | 0.164 | 0.262 | 0.186 (4) | 0.262 (5) |
| Taylor | 0.230 (9†) | 0.520 (63) | 0.338 | 0.494 (12) | 0.335 | 0.488 (6) |

† Number of days in which soil water content in one of the soil depths was below the wilting point.

### Supplementary Material B: Relationship between dryland cotton yield and adjustments in $\pi_o$ and LDMC

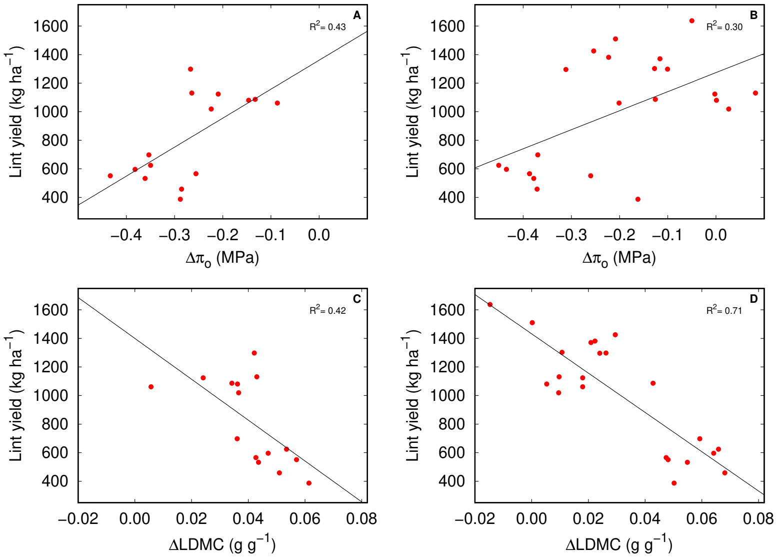


Figure B1: Dryland lint yield of selected cotton varieties as predicted by the adjustments in leaf osmotic potential at full turgor ($\Delta\pi_o$) and leaf dry matter content (ΔLDMC) at the early bloom (A, C) and peak bloom (B, D) stages. The adjustments in leaf osmotic potential and leaf dry matter content are defined as differences in measured values at the dryland site and at the irrigated site for the same varieties in the same year from 2020 to 2022.

### Supplementary Material C: Drought Tolerance Ranking of Cotton Varieties

#### C1 Data for drought tolerance ranking

As explained in the main text, $p = 17$ varieties in $m = 6$ different seasons are used for the statistical ranking problem. In total, we have $n = 280$ observations of LDMC, details of which are given in Table C1.

Table C1: Number of replicates for LDMC measurement by cotton variety and season ('early' denotes the early bloom season and 'peak' denotes the peak bloom season). For each variety measured in a season, 2 or 3 replicates are taken at the dryland site and 3 replicates at the irrigated site. The $\pi_o$ measurement is also obtained for all replicates, except that there are no dryland measurements for $\pi_o$ in the peak bloom season of year 2021 due to unforeseen circumstances.

| | 2020 | | 2021 | | 2022 | |
|---|---|---|---|---|---|---|
| Variety | early | peak | early | peak | early | peak |
| DG 3456 | 0 | 0 | 0 | 0 | 6 | 6 |
| DG 3555 | 0 | 0 | 0 | 0 | 6 | 6 |
| DP 1646 | 6 | 6 | 6 | 6 | 0 | 0 |
| DP 2012 | 0 | 0 | 0 | 0 | 6 | 6 |
| DP 2020 | 6 | 6 | 6 | 6 | 6 | 6 |
| H 959 | 0 | 0 | 6 | 6 | 0 | 0 |
| NG 4098 | 6 | 6 | 0 | 0 | 0 | 0 |
| NG 4190 | 0 | 0 | 0 | 0 | 6 | 6 |
| NG 4936 | 6 | 6 | 5 | 5 | 0 | 0 |
| NG 5150 | 0 | 0 | 5 | 5 | 0 | 0 |
| PHY 332 | 0 | 0 | 5 | 5 | 6 | 6 |
| PHY 400 | 6 | 6 | 5 | 5 | 6 | 6 |
| PHY 480 | 6 | 6 | 0 | 0 | 0 | 0 |
| ST 4550 | 6 | 6 | 0 | 0 | 0 | 0 |
| ST 4595 | 0 | 0 | 0 | 0 | 6 | 6 |
| ST 4990 | 6 | 6 | 0 | 0 | 0 | 0 |
| ST 5707 | 6 | 6 | 0 | 0 | 0 | 0 |

#### C2 Bayesian joint modeling of leaf dry matter content and leaf osmotic potential

Recall that we denote the set of $p$ cotton varieties by $\mathcal{V}$ and the set of $m$ seasons by $\mathcal{S}$. For the $k$-th observation ($k = 1, 2, \ldots, n$), $Y^{\mathrm{D}}(k)$ denotes the standardized LDMC measurement, $Y^{\pi}(k)$ denotes the standardized *negative* $\pi_o$ measurement, $\mathrm{vat}(k) \in \mathcal{V}$ denotes the cotton variety, $\mathrm{sea}(k) \in \mathcal{S}$ denotes the season, and $\mathrm{env}(k) \in \{0, 1\}$ denotes the environmental condition (1 for dryland). We assume that

$$Y^{\mathrm{D}}(k) = \mu^{\mathrm{D}}_{\mathrm{vat}(k)} + a^{\mathrm{D}}_{\mathrm{vat}(k)} \mathrm{env}(k) + c^{\mathrm{D}}_{\mathrm{sea}(k)} + e^{\mathrm{D}}_k, \quad e^{\mathrm{D}}_k \overset{\mathrm{ind}}{\sim} N(0, \sigma^2_{\mathrm{D}}), \tag{1}$$

$$Y^{\pi}(k) = \mu^{\pi}_{\mathrm{vat}(k)} + a^{\pi}_{\mathrm{vat}(k)} \mathrm{env}(k) + c^{\pi}_{\mathrm{sea}(k)} + e^{\pi}_k, \quad e^{\pi}_k \overset{\mathrm{ind}}{\sim} N(0, \sigma^2_{\pi}), \tag{2}$$

where $e_k^{\mathrm{D}}, e_k^{\pi}$ are independent Gaussian errors, $\{\mu_v^{\mathrm{D}}, \mu_v^{\pi}, a_v^{\mathrm{D}}, a_v^{\pi} : v \in \mathcal{V}\}$ and $\{c_s^{\mathrm{D}}, c_s^{\pi} : s \in \mathcal{S}\}$ are unknown parameters of interest, and $\sigma_{\mathrm{D}}^2, \sigma_{\pi}^2$ are unknown hyperparameters. We use $\boldsymbol{\mu}^{\mathrm{D}}$ to denote the vector that contains $p$ elements $\{\mu_v^{\mathrm{D}} : v \in \mathcal{V}\}$, assuming an implicit one-to-one mapping from $\mathcal{V}$ to $\{1, 2, \ldots, p\}$; $\boldsymbol{\mu}^{\pi}, \boldsymbol{a}^{\mathrm{D}}, \boldsymbol{a}^{\pi}$, $\boldsymbol{c}^{\mathrm{D}}, \boldsymbol{c}^{\pi}$ are defined similarly, but note that $\boldsymbol{\mu}^{\mathrm{D}}, \boldsymbol{\mu}^{\pi}, \boldsymbol{a}^{\mathrm{D}}, \boldsymbol{a}^{\pi} \in \mathbb{R}^p$ while $\boldsymbol{c}^{\mathrm{D}}, \boldsymbol{c}^{\pi} \in \mathbb{R}^m$. The parameter $\boldsymbol{a}^{\mathrm{D}}$ (resp. $\boldsymbol{a}^{\pi}$) characterizes the difference in LDMC (resp. $\pi_o$) measurement for the same cotton variety under dryland and irrigated conditions. If the leaf of a cotton variety $v$ has better drought tolerance, then both $a_v^{\mathrm{D}}$ and $a_v^{\pi}$ tend to be larger.

The main idea of Bayesian statistical modeling is to first specify a prior distribution for unknown model parameters and then update it using the data likelihood, which yields the posterior distribution of parameters. Compared to standard Bayesian linear regression models, the key novelty of our method is that our prior forces the drought tolerance parameters $\{a_v^{\mathrm{D}} : v \in \mathcal{V}\}$ and $\{a_v^{\pi} : v \in \mathcal{V}\}$ to have the same ranking. Further, our prior encodes the constraint that $a_v^{\mathrm{D}} \geqslant 0, a_v^{\pi} \geqslant 0$ for all $p$ varieties. In words, our prior assumes that the observed data is generated from the following process.

Step 1. Draw $\sigma_{\mathrm{D}}^2, \sigma_{\pi}^2, \boldsymbol{\mu}^{\mathrm{D}}, \boldsymbol{\mu}^{\pi}, \boldsymbol{c}^{\mathrm{D}}, \boldsymbol{c}^{\pi}$ from the prior distributions given in (7), (8) and (9).

Step 2. Draw an ordering of the $p$ cotton varieties, denoted by $\tau$, from the prior distribution given in (4).

Step 3. Draw $\boldsymbol{a}^{\mathrm{D}}$ and $\boldsymbol{a}^{\pi}$ from some normal prior distributions under the constraints that (i) $a_v^{\mathrm{D}} \geqslant 0, a_v^{\pi} \geqslant 0$ for all $v \in \mathcal{V}$, and (ii) both $\boldsymbol{a}^{\mathrm{D}}$ and $\boldsymbol{a}^{\pi}$ are consistent with the ordering $\tau$. See (5) and (6).

Step 4. Generate observations according to (1) and (2).

Formally, let $\mathbb{S}_{\mathrm{vat}}$ denote the set of all possible total orderings of the elements in $\mathcal{V}$, which has $p!$ elements. Given $\tau \in \mathbb{S}_{\mathrm{vat}}$, $\tau(i) = v$ means that the variety $v$ has drought tolerance ranking equal to $i$ (a larger $i$ indicates better drought tolerance). For each $\tau \in \mathbb{S}_{\mathrm{vat}}$, define

$$\mathbb{R}_{+,\tau}^p = \{\boldsymbol{a} \in \mathbb{R}^p : 0 \leqslant a_{\tau(1)} \leqslant a_{\tau(2)} \leqslant \cdots \leqslant a_{\tau(p)}\}, \tag{3}$$

which is the set of all possible values of $\boldsymbol{a}$ consistent with ordering $\tau$. Let $N(m, V, S)$ denote the normal distribution with mean $m$ and covariance matrix $V$ constrained to be in the set $S$. We use the following hierarchical prior for $\boldsymbol{a}^{\mathrm{D}}, \boldsymbol{a}^{\pi}$:

$$\tau \sim \mathrm{Unif}(\mathbb{S}_{\mathrm{vat}}), \tag{4}$$
$$\boldsymbol{a}^{\mathrm{D}} \mid \tau \sim N(\boldsymbol{m}^{\mathrm{D}}, \nu_a^{\mathrm{D}} \boldsymbol{I}_p, \mathbb{R}_{+,\tau}^p), \quad v \in \mathcal{V}, \tag{5}$$
$$\boldsymbol{a}^{\pi} \mid \tau \sim N(\boldsymbol{m}^{\pi}, \nu_a^{\pi} \boldsymbol{I}_p, \mathbb{R}_{+,\tau}^p), \quad v \in \mathcal{V}, \tag{6}$$

where $\boldsymbol{m}^{\mathrm{D}}, \boldsymbol{m}^{\pi} \in \mathbb{R}^p, \nu_a^{\mathrm{D}}, \nu_a^{\pi} > 0$ are hyperparameters, $\boldsymbol{I}_p$ denotes the $p \times p$ identity matrix, and $\mathrm{Unif}(\mathbb{S}_{\mathrm{vat}})$ denotes the uniform distribution on $\mathbb{S}_{\mathrm{vat}}$. For the other parameters, we simply use the standard normal-inverse-gamma prior distributions:

$$\sigma_{\mathrm{D}}^2, \sigma_{\pi}^2 \overset{\mathrm{ind}}{\sim} \text{Inverse-gamma}(\kappa_1, \kappa_2), \tag{7}$$
$$\boldsymbol{\mu}^{\mathrm{D}} \sim N(0, \nu_\mu^{\mathrm{D}} \boldsymbol{I}_p), \qquad \boldsymbol{\mu}^{\pi} \sim N(0, \nu_\mu^{\pi} \boldsymbol{I}_p), \tag{8}$$
$$\boldsymbol{c}^{\mathrm{D}} \sim N(0, \nu_c^{\mathrm{D}} \boldsymbol{I}_g), \qquad \boldsymbol{c}^{\pi} \sim N(0, \nu_c^{\pi} \boldsymbol{I}_g). \tag{9}$$

For the hyperparameters involved in the prior normal distributions, we choose their values by first fitting a linear regression model with response $Y^{\mathrm{D}}$ and that with response $Y^{\pi}$ separately using least squares. According to the estimated regression coefficients, we decided to use

$$\begin{aligned}\nu_\mu^{\mathrm{D}} &= \nu_\mu^{\pi} = 0.5,\\ \nu_a^{\mathrm{D}} &= \nu_a^{\pi} = 0.5,\\ \nu_c^{\mathrm{D}} &= \nu_c^{\pi} = 1,\\ \boldsymbol{m}^{\mathrm{D}} &= \boldsymbol{m}^{\mathrm{D}} = \mathbf{1},\end{aligned}$$

where $\mathbf{1}$ denotes a vector of which all elements equal 1. The variance hyperparameters are larger (approximately double) the variances of least-squares regression coefficient estimates, since we do not want to make the prior too informative. Finally, we let $\kappa_1 = \kappa_2 = 0.5$, which is a standard non-informative choice in Bayesian linear models.

### C3 Conditional posterior distributions

Due to the use of conjugate normal-inverse-gamma prior, the full conditional posterior distributions of $\boldsymbol{\mu}^{\mathrm{D}}, \boldsymbol{\mu}^{\pi}, \boldsymbol{c}^{\mathrm{D}}, \boldsymbol{c}^{\pi}, \sigma^2_{\mathrm{D}}, \sigma^2_{\pi}$ can be obtained straightforwardly. For example, consider $\boldsymbol{\mu}^{\mathrm{D}}$. For each $k = 1, \dots, n$, define

$$\hat{\mu}^{\mathrm{D}}(k) = Y^{\mathrm{D}}(k) - a^{\mathrm{D}}_{\mathrm{vat}(k)}\mathrm{env}(k) - c^{\mathrm{D}}_{\mathrm{sea}(k)},$$

which can be evaluated if all the parameters except $\boldsymbol{\mu}^{\mathrm{D}}$ are given. Let $\mathcal{I}_v^{\mathrm{D}} = \{1 \leqslant k \leqslant n \colon \mathrm{vat}(k) = v, Y^{\mathrm{D}}(k) \text{ is observed}\}$ denote the index set of all records for which the observed variety is $v$ and LDMC measurement is not missing,[1] and let $n_v^{\mathrm{D}}$ denote the number of elements in $\mathcal{I}_v^{\mathrm{D}}$. The full conditional posterior distribution of $\mu_v^{\mathrm{D}}$ is still a normal distribution:

$$\mu_v^{\mathrm{D}} \mid (Y^{\mathrm{D}}(k))_k, \boldsymbol{a}^{\mathrm{D}}, \boldsymbol{c}^{\mathrm{D}}, \sigma^2_{\mathrm{D}} \sim N\left(\frac{\sum_{k\in\mathcal{I}_v^{\mathrm{D}}} \hat{\mu}^{\mathrm{D}}(k)/\sigma^2_{\mathrm{D}}}{1/\nu_\mu^{\mathrm{D}} + n_v^{\mathrm{D}}/\sigma^2_{\mathrm{D}}}, \frac{1}{1/\nu_\mu^{\mathrm{D}} + n_v^{\mathrm{D}}/\sigma^2_{\mathrm{D}}}\right).$$

The full conditional posterior distributions of $\boldsymbol{\mu}^{\pi}, \boldsymbol{c}^{\mathrm{D}}, \boldsymbol{c}^{\pi}$ can be calculated similarly. For $\sigma^2_{\mathrm{D}}$, its full conditional posterior distribution is still the inverse-gamma distribution:

$$\sigma^2_{\mathrm{D}} \mid (Y^{\mathrm{D}}(k))_k, \boldsymbol{a}^{\mathrm{D}}, \boldsymbol{c}^{\mathrm{D}}, \mu_v^{\mathrm{D}} \sim \text{Inverse-gamma}\left(\kappa_1 + \frac{n^{\mathrm{D}}}{2}, \kappa_2 + \frac{1}{2}\sum_{k\in\mathcal{I}^{\mathrm{D}}} (\hat{e}_k^{\mathrm{D}})^2\right),$$

where $n^{\mathrm{D}} = \sum_{v\in\mathcal{V}} n_v^{\mathrm{D}}$, $\mathcal{I}^{\mathrm{D}} = \cup_{v\in\mathcal{V}}\mathcal{I}_v^{\mathrm{D}}$, and

$$\hat{e}_k^{\mathrm{D}} = Y^{\mathrm{D}}(k) - \mu^{\mathrm{D}}_{\mathrm{vat}(k)} - a^{\mathrm{D}}_{\mathrm{vat}(k)}\mathrm{env}(k) - c^{\mathrm{D}}_{\mathrm{sea}(k)}.$$

Note that since there are no missing LDMC measurments, we have $n^{\mathrm{D}} = n$ and $\mathcal{I}^{\mathrm{D}} = \{1, 2, \dots, n\}$. The full conditional posterior distribution of $\sigma^2_{\pi}$ can be obtained analogously.

Consider $a_v^{\mathrm{D}}$ for some $v \in \mathcal{V}$. Given $\tau$, $\boldsymbol{a}^{\mathrm{D}}_{-v}$ and other parameters, the full conditional posterior distribution of $a_v^{\mathrm{D}}$ is still a truncated normal distribution. For each $k$ such that $\mathrm{env}(k) = 1$, define

$$\hat{a}^{\mathrm{D}}(k) = Y^{\mathrm{D}}(k) - \mu^{\mathrm{D}}_{\mathrm{vat}(k)} - c^{\mathrm{D}}_{\mathrm{sea}(k)}.$$

[1] For all the $n$ records, LDMC is observed; thus, the condition "$Y^{\mathrm{D}}(k)$ is observed" can be dropped in the definition of $\mathcal{I}_v^{\mathrm{D}}$. But note that there are missing observations for $Y^{\pi}$.

Let $\mathcal{I}^{\mathrm{D}}_{v,1} = \{1 \leqslant k \leqslant n\colon \mathrm{vat}(k) = v, \mathrm{env}(k) = 1,\, Y^{\mathrm{D}}(k) \text{ is observed}\}$, and $n^{\mathrm{D}}_{v,1}$ denote the number of elements in $\mathcal{I}^{\mathrm{D}}_{v,1}$. We have

$$
\begin{aligned}
& a^{\mathrm{D}}_v \mid (Y^{\mathrm{D}}(k))_k, \tau, \boldsymbol{a}^{\mathrm{D}}_{-v}, \boldsymbol{\mu}^{\mathrm{D}}, \boldsymbol{c}^{\mathrm{D}}, \sigma^2_{\mathrm{D}} \\
& \sim N\left( \frac{\sum_{k \in \mathcal{I}^{\mathrm{D}}_{v,1}} \hat{a}^{\mathrm{D}}(k)/\sigma^2_{\mathrm{D}}}{1/\nu^{\mathrm{D}}_a + n^{\mathrm{D}}_{v,1}/\sigma^2_{\mathrm{D}}},\ \frac{1}{1/\nu^{\mathrm{D}}_a + n^{\mathrm{D}}_{v,1}/\sigma^2_{\mathrm{D}}},\ (a_0, a_1) \right), \\
& \text{where } a_0 = a^{\mathrm{D}}_{\tau(\tau^{-1}(v)-1)}, \text{ and } a_1 = a^{\mathrm{D}}_{\tau(\tau^{-1}(v)+1)}.
\end{aligned}
$$

If $\tau^{-1}(v) = 1$, we set $a_0 = -\infty$; if $\tau^{-1}(v) = p$, we set $a_1 = +\infty$.

### C4 Markov chain Monte Carlo sampling

We use Markov chain Monte Carlo (MCMC) sampling to learn the joint posterior distribution of the parameters $\tau$, $\boldsymbol{\mu}^{\mathrm{D}}, \boldsymbol{\mu}^{\pi}, \boldsymbol{a}^{\mathrm{D}}, \boldsymbol{a}^{\pi}, \boldsymbol{c}^{\mathrm{D}}, \boldsymbol{c}^{\pi}, \sigma^2_{\mathrm{D}}, \sigma^2_{\pi}$. Due to the use of conjugate prior distributions, we are able to build an efficient random-scan Metropolis-within-Gibbs algorithm. All parameters except $\tau$ can be updated from their full conditional distributions (Gibbs update), and we use an acceptance-rejection step to update the ordering $\tau$ (Metropolis update). The algorithm is detailed below.

**Algorithm 1.** Choose positive constants $q_\mu, q_a, q_c, q_\sigma, q_\tau$ such that $q_\mu + q_a + q_c + q_\sigma + q_\tau = 1$. In each MCMC iteration, do the following.

- Generate `Type` from the following distribution:

$$
\begin{gathered}
P(\texttt{Type} = 1) = q_\mu, \quad P(\texttt{Type} = 2) = q_a, \quad P(\texttt{Type} = 3) = q_c, \\
P(\texttt{Type} = 4) = q_\sigma, \quad P(\texttt{Type} = 5) = q_\tau.
\end{gathered}
$$

- If `Type` $= 1$, draw $v$ from $\mathcal{V}$ randomly and update $\mu^{\mathrm{D}}_v, \mu^{\pi}_v$ from their full conditional posterior distributions.
- If `Type` $= 2$, draw $v$ from $\mathcal{V}$ randomly and update $a^{\mathrm{D}}_v, a^{\pi}_v$ from their full conditional posterior distributions.
- If `Type` $= 3$, draw $s$ from $\mathcal{S}$ randomly and update $c^{\mathrm{D}}_s, c^{\pi}_s$ from their full conditional posterior distributions.
- If `Type` $= 4$, update $\sigma^2_{\mathrm{D}}, \sigma^2_{\pi}$ from their full conditional posterior distributions.
- If `Type` $= 5$, draw $i$ from $1, 2, \dots, p-1$ randomly. Let $\tilde{\tau}$ be obtained from $\tilde{\tau}$ by swapping the $i$-th and $(i+1)$-th elements of $\tau$, let $\tilde{\boldsymbol{a}}^{\mathrm{D}}$ be obtained from $\boldsymbol{a}^{\mathrm{D}}$ by swapping $a^{\mathrm{D}}_{\tau(i)}$ and $a^{\mathrm{D}}_{\tau(i+1)}$, and let $\tilde{\boldsymbol{a}}^{\pi}$ be obtained from $\boldsymbol{a}^{\pi}$ by swapping $a^{\pi}_{\tau(i)}$ and $a^{\pi}_{\tau(i+1)}$. Let $\boldsymbol{\theta} = (\boldsymbol{\mu}^{\mathrm{D}}, \boldsymbol{\mu}^{\pi}, \boldsymbol{c}^{\mathrm{D}}, \boldsymbol{c}^{\pi}, \sigma^2_{\mathrm{D}}, \sigma^2_{\pi})$ denote all the other parameters, and $L$ denote the likelihood function (see below). With probability

$$
\min\left\{1, \frac{L(\tilde{\tau}, \tilde{\boldsymbol{a}}^{\mathrm{D}}, \tilde{\boldsymbol{a}}^{\pi}, \boldsymbol{\theta})}{L(\tau, \boldsymbol{a}^{\mathrm{D}}, \boldsymbol{a}^{\pi}, \boldsymbol{\theta})}\right\}, \tag{10}
$$

  we accept the proposal and update $(\tau, \boldsymbol{a}^{\mathrm{D}}, \boldsymbol{a}^{\pi})$ to $(\tilde{\tau}, \tilde{\boldsymbol{a}}^{\mathrm{D}}, \tilde{\boldsymbol{a}}^{\pi})$. If the proposal is rejected, we do nothing.

In the above algorithm, the data likelihood can be calculated by

$$
\begin{aligned}
&L(\tau, \boldsymbol{a}^{\mathrm{D}}, \boldsymbol{a}^{\pi}, \boldsymbol{\mu}^{\mathrm{D}}, \boldsymbol{\mu}^{\pi}, \boldsymbol{c}^{\mathrm{D}}, \boldsymbol{c}^{\pi}, \sigma^2_{\mathrm{D}}, \sigma^2_{\pi}) \\
= &\left\{ \prod_{k \in \mathcal{I}^{\mathrm{D}}} \phi \left( \frac{Y^{\mathrm{D}}(k) - \mu^{\mathrm{D}}_{\mathrm{vat}(k)} - a^{\mathrm{D}}_{\mathrm{vat}(k)} \mathrm{env}(k) - c^{\mathrm{D}}_{\mathrm{sea}(k)}}{\sigma_{\mathrm{D}}} \right) \right\} \times \\
&\left\{ \prod_{k \in \mathcal{I}^{\pi}} \phi \left( \frac{Y^{\pi}(k) - \mu^{\pi}_{\mathrm{vat}(k)} - a^{\pi}_{\mathrm{vat}(k)} \mathrm{env}(k) - c^{\pi}_{\mathrm{sea}(k)}}{\sigma_{\pi}} \right) \right\},
\end{aligned}
$$

where $\phi$ denotes the density function of the standard normal distribution with mean zero and variance one. Note that when evaluating (10), we only need to calculate the likelihood for those observations that involve either variety $\tau(i)$ or variety $\tau(i+1)$. The standard MCMC theory implies that Algorithm 1 constructs a Markov chain which converges to the joint posterior distribution of $\tau, \boldsymbol{a}^{\mathrm{D}}, \boldsymbol{a}^{\pi}, \boldsymbol{\mu}^{\mathrm{D}}, \boldsymbol{\mu}^{\pi}, \boldsymbol{c}^{\mathrm{D}}, \boldsymbol{c}^{\pi}, \sigma^2_{\mathrm{D}}, \sigma^2_{\pi}$. We run Algorithm 1 for $10^6$ iterations with half of them as burn-in, which only takes a few minutes on a personal laptop. Trace plots suggest that the algorithm has approximately converged after $10^4$ iterations.

#### C5 Results

Figures C1 and C2 visualize the posterior mean estimates (presented in Table 1) and their associated 50% credible intervals. To demonstrate the advantage of joint modeling of LDMC and $\pi_o$, we also conduct separate analysis of LDMC and $\pi_o$ using the same Bayesian linear model. The latent variable $\tau$ is not needed in the separate analysis, and all model parameters can be efficiently sampled via Gibbs updates. Prior hyperparameters and MCMC setting of the separate analysis are identical to those used in the joint analysis. The posterior distributions of $\boldsymbol{a}^{\mathrm{D}}$ and $\boldsymbol{a}^{\pi}$ are shown in the second row of Figure C2, and it can be seen that the estimates have significantly larger variation than in the joint analysis. This is because the sample size of our data is relatively small, and thus combining information from LDMC and $\pi_o$ can significantly increase the precision of posterior estimates.

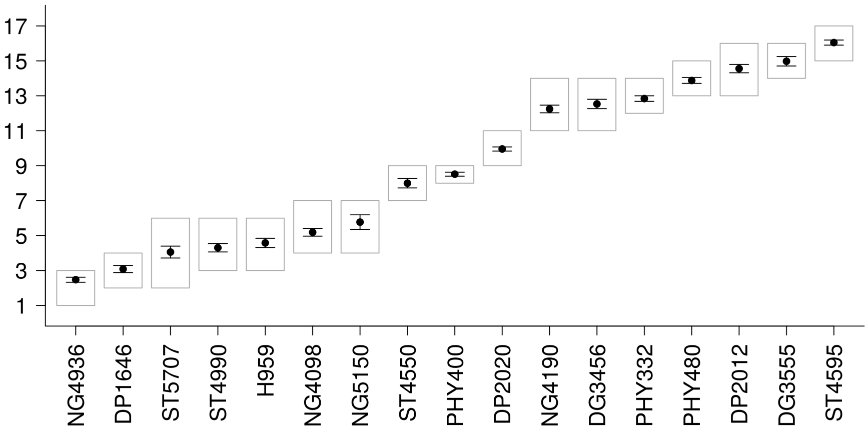


Figure C1: Drought tolerance ranking of 17 cotton varieties via joint Bayesian analysis of LDMC and $\pi_o$. The solid dot represents the posterior mean value of the ranking estimated from the MCMC output, and the black error bars indicate $\pm$3SE. The standard error of the posterior mean is estimated by splitting the MCMC samples into 10 batches. The gray box represents the 50% credible interval, i.e., the 25% and 75% quantiles of MCMC samples. The 17 varieties are ordered according to the posterior mean ranking. A larger ranking indicates a higher capacity of drought tolerance.

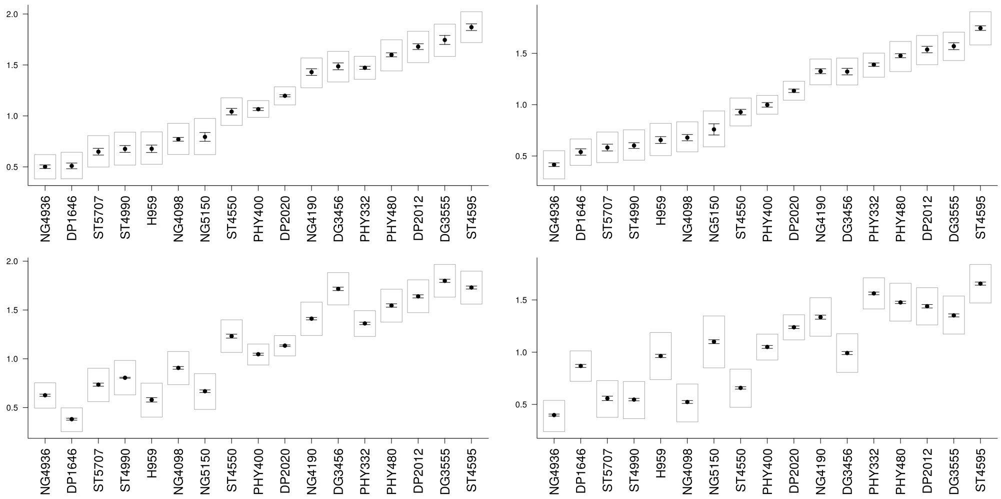

Figure C2: Estimates of $\boldsymbol{a}^{\mathrm{D}}$ and $\boldsymbol{a}^{\pi}$ from the MCMC output. Top left: $\boldsymbol{a}^{\mathrm{D}}$ from joint analysis; top right: $\boldsymbol{a}^{\pi}$ from joint analysis; bottom left: $\boldsymbol{a}^{\mathrm{D}}$ from separate analysis; bottom right: $\boldsymbol{a}^{\pi}$ from separate analysis. As in Figure C1, the solid dot represents the posterior mean, the error bars represent $\pm$3SE, and the gray box represents the 50% credible interval. In all the 4 panels, varieties are ordered in the same way as in Figure C1.

**Supplementary Material D.** Table D1. Probability values of the ANOVA results for variables of leaf gas exchange and water relations measurements presented in Figure 7 of the main text. The effects of the fixed factors time (T), irrigation treatments (D), genotypes (G) and their interactions (T x I, T x G, I x G and T x I x G) are considered, and the variables include foliar gas exchange (net photosynthetic rate (A), carboxylation efficiency ($A/C_i$), stomatal conductance ($g_s$), intrinsic water use efficiency ($A/g_s$), instantaneous water use efficiency (A/E), maximum photochemical efficiency of PSII ($F_v/F_m$), thylakoid membrane damage ($F_0/F_m$), leaf water potential ($\psi_L$), leaf temperature ($T_{leaf}$), and canopy temperature ($T_{canopy}$).

| Variables | | | | | | | |
|---|---|---|---|---|---|---|---|
| Traits | Time (T) | Irrigation (I) | Genotypes (G) | T x I | T x G | I x G | Tx I x G |
| A | <0.001 | <0.001 | <0.001 | <0.001 | <0.001 | <0.001 | <0.01 |
| A/Ci | <0.001 | <0.001 | <0.05 | <0.001 | <0.001 | <0.001 | 0.073 |
| gs | <0.001 | <0.001 | <0.001 | <0.001 | <0.001 | <0.001 | <0.01 |
| A/gs | <0.001 | <0.001 | <0.001 | <0.001 | <0.001 | <0.001 | <0.005 |
| A/E | <0.001 | <0.05 | <0.005 | 0.091 | <0.05 | <0.05 | <0.001 |
| Fv/Fm | 0.370 | 0.320 | 0.371 | 0.372 | 0.412 | 0.372 | 0.412 |
| F0/Fm | <0.001 | 0.279 | <0.001 | 0.113 | 0.487 | 0.961 | 0.324 |
| ψL | <0.001 | <0.001 | <0.001 | <0.001 | 0.393 | <0.001 | 0.478 |
| Tleaf | <0.001 | <0.001 | 0.122 | <0.001 | <0.05 | 0.122 | <0.001 |
| Tcanopy | <0.001 | <0.001 | <0.001 | <0.001 | <0.001 | <0.001 | <0.001 |
